# A PCNA-K164R mutation impinges on origin activation and mitotic DNA synthesis

**DOI:** 10.1101/2020.06.25.172361

**Authors:** Wendy Leung, Ryan M. Baxley, Tanay Thakar, Ya-Chu Chang, Colette B. Rogers, Liangjun Wang, Wesley Durrett, Anika Tella, George-Lucian Moldovan, Naoko Shima, Anja-Katrin Bielinsky

## Abstract

Ubiquitination of the replication clamp proliferating cell nuclear antigen (PCNA) at the conserved residue lysine 164 (K164) occurs during normal S phase progression and increases after DNA damage induced replication stress. PCNA-K164 ubiquitination is critical for Okazaki fragment maturation and the activation of DNA damage tolerance pathways. Moreover, ubiquitinated PCNA operates in a fork protection pathway parallel to BRCA-RAD51. Whether PCNA ubiquitination regulates other genome maintenance mechanisms is unclear. Utilizing *PCNA^K164R^* cells generated by CRISPR-Cas9, we demonstrate that this mutation causes DNA replication defects that impact origin activation. *PCNA^K164R^* cells accumulate single-stranded DNA gaps during replication that persist throughout mitosis due to compromised mitotic DNA synthesis (MiDAS). We uncover a novel role for PCNA-K164 ubiquitination in regulating FANCD2 to initiate MiDAS. Persistent gaps hence interfere with MCM2-7 double hexamer loading in the subsequent G1 phase. Our findings demonstrate that the impact of PCNA^K164-Ub^ is not limited to S/G2 phases but extends to mitosis and G1 phase.

**SUMMARY:** PCNA-K164 ubiquitination promotes DNA gap filling during S/G2 phases of the cell cycle. This study identifies a novel role for K164 ubiquitination in replication dynamics and mitotic DNA synthesis and thus provides new insight into the players involved in counteracting under-replication.

## INTRODUCTION

Maintenance of genome integrity is an intricate process that requires an extended protein network and coordination of multiple cellular pathways (Barnes and Eckert, 2017). Proper DNA replication is essential to genome maintenance and ensures precise duplication without leaving sequences un-replicated or replicated more than once. Replication can be divided into three phases: origin licensing, origin firing and DNA synthesis (Fragkos et al., 2015). Origin licensing occurs during late mitosis and G1 phase where cell division cycle (CDC) protein 6 (CDC6) and chromatin licensing and DNA replication factor 1 (CDT1) direct the loading of the minichromosome maintenance (MCM) proteins 2-7 (MCM2-7) in a double hexameric complex onto DNA (Donovan et al., 1997; Seki and Diffley, 2000; Remus et al., 2009; Evrin et al., 2009; Siddiqui et al., 2013). As the cell transitions from G1 to S phase, the helicase co-activators CDC45 and the go-ichi-ni-san (GINS) complex are recruited to form the CDC45-MCM2-7-GINS (CMG) helicase (Bochman and Schwacha, 2008; Moyer et al., 2006). To initiate DNA synthesis, a pair of CMG complexes is activated and the recruitment of MCM10, proliferating cell nuclear antigen (PCNA), and DNA polymerases α, ε, and δ occurs to catalyze bidirectional DNA replication (Rivera-Mulia and Gilbert, 2016; Burgers and Kunkel, 2017; Baxley and Bielinsky, 2017; Langston and O’Donnell, 2019).

Endogenous and exogenous sources of DNA damage can impede replication and generate aberrant fork structures that can give rise to chromosomal alterations (Macheret and Halazonetis, 2015; Técher et al., 2017; Tubbs and Nussenzweig, 2017; Primo and Teixeira, 2020). In response to these challenges, PCNA is ubiquitinated at the conserved lysine (K) residue 164, activating DNA damage tolerance (DDT) pathways (Friedberg, 2005; Chang and Cimprich, 2009; Ghosal and Chen, 2013; Leung et al., 2018). DDT pathways are categorized as either error-prone or error-free. Mono-ubiquitination at K164 by the E2-E3 radiation sensitive (RAD) protein 6 and 18 (RAD6-RAD18) complex activates the error-prone translesion synthesis (TLS) pathway (Hoege et al., 2002; Davies et al., 2008). TLS is catalyzed by specialized low-fidelity DNA polymerases to directly bypass DNA lesions (Shcherbakova and Fijalkowska, 2006; Lehmann et al., 2007; Sale et al., 2012). Mono-ubiquitinated PCNA can be further modified by K63-linked poly-ubiquitin chains by the E2 ubiquitin conjugating enzyme UBC13-MMS21 and two E3 ligases, helicase-like transcription factor (HLTF) or SNF2 histone linker PHD RING helicase (SHPRH), driving error-free template switching (TS) (Broomfield et al., 1998; Hofmann and Pickart, 1999; Ulrich, 2000; Brusky et al., 2000; Stelter and Ulrich, 2003; Branzei et al., 2004; Unk et al., 2006, 2008; Motegi et al., 2006, 2008). TS utilizes a recombination-like mechanism by which the nascent DNA of the sister chromatid serves as a template for replication (Vanoli et al., 2010; Minca and Kowalski, 2010; Branzei, 2011; Gonzalez-Huici et al., 2014; Fumasoni et al., 2015; Branzei and Szakal, 2016). To facilitate this switch, K63-linked ubiquitin chains recruit the DNA translocase, zinc finger RAN-binding domain containing protein 3 (ZRANB3), to reverse the fork (Ciccia et al., 2012; Weston et al., 2012; Yuan et al., 2012; Badu-Nkansah et al., 2016; Vujanovic et al., 2017). Given the importance of PCNA-K164 ubiquitination, it is not surprising that mutating this residue renders cells hypersensitive to DNA damage (Arakawa et al., 2006; Edmunds et al., 2008; Hendel et al., 2011; Niimi et al., 2008; Qin et al., 2013). However, how this modification and its associated DDT pathways function in maintaining human genome stability under unperturbed conditions is still not understood.

Regions of the genome that may be particularly reliant on DDT pathways include late-replicating common fragile sites (CFS) (Le Beau et al., 1998). Failed replication at CFS can be caused by an insufficient number of licensed origins, inefficient origin firing, and failure to activate dormant origins (Palakodeti et al., 2009; Letessier et al., 2011; Ozeri-Galai et al., 2011). Insufficient origin licensing can increase late replication intermediates (LRIs) at CFS as well as other loci, leading to the formation of Fanconi Anemia (FA) group D2 protein (FANCD2) foci during G2/M phase (Kawabata et al., 2011; Chan et al., 2009). Such LRIs can be resolved through the formation of ultra-fine bridges in anaphase and/or p53-binding protein 1 (53BP1) nuclear bodies (NBs) during the subsequent G1- or S phase. A recently discovered mechanism to resolve LRIs is mitotic DNA synthesis (MiDAS) (Minocherhomji et al., 2015; Bhowmick et al., 2016). MiDAS employs a break-induced replication (BIR)-like process to resolve LRIs and to ensure proper chromosome segregation (Minocherhomji et al., 2015; Bhowmick et al., 2016; Graber-Feesl et al., 2019). While MiDAS in human cancer cells depends on RAD52 (Bhowmick et al., 2016), we recently demonstrated that this RAD52-driven mechanism is absent and instead relies on FANCD2 in nontransformed human cells (Graber-Feesl et al., 2019). Whether PCNA ubiquitination plays any role in MiDAS has not been explored, although these pathways cooperate in DNA crosslink repair (Kim and D’Andrea, 2012).

We have recently shown that PCNA-K164 ubiquitination acts in parallel to breast cancer (BRCA) predisposition genes 1 and 2 and the FA pathway to prevent nascent DNA degradation by DNA replication ATP-dependent helicase/nuclease, DNA2 (Thakar et al., 2020). Although fork reversal is a mechanism to restart replication, reversed forks are often subjected to nucleolytic degradation. Loading of RAD51 onto nascent DNA at stalled forks by the BRCA1/2-FA pathway prevents MRE11 nuclease-dependent degradation (Schlacher et al., 2011, 2012; Couch et al., 2013; Tirman et al., 2021). When this pathway is defective, PCNA-K164 ubiquitination becomes critical to promote efficient Okazaki fragment (OF) ligation, allowing for proper PCNA unloading and nucleosome deposition, thereby preventing DNA2 nuclease-dependent degradation (Thakar et al., 2020). Thus, both the BRCA1/2-FA and PCNA-K164 ubiquitination pathways function in fork protection. Additionally, Nayak and co-workers revealed that activation of TLS polymerases prevents replication fork slowing and remodeling, thereby suppressing ssDNA gap formation and promoting cancer cell fitness. Impairing TLS, which acts predominantly in G2 phase (Tirman et al., 2021), disrupts DNA replication and synergizes with gap-inducing therapies, highlighting the importance of replication gaps as a cancer vulnerability (Nayak et al., 2020).

In this study, we demonstrate that PCNA-K164 ubiquitination is critical for accurate DNA replication and genome stability in human cells. Loss of PCNA-K164 ubiquitination in 293T and hTERT immortalized RPE-1 (referred to subsequently as RPE-1) cells leads to decreased origin licensing and firing, and impaired DNA synthesis under unperturbed conditions. These defects generate ssDNA gaps that are not resolved during mitosis. Compromised MiDAS in *PCNA^K164R^* mutants is directly linked to decreased FANCD2 mono-ubiquitination, which ultimately leads to anaphase aberrations and the accumulation of 53BP1 NBs. Taken together, our data show that PCNA ubiquitination at K164 is not only important for progressive S phase DNA synthesis but promotes efficient origin licensing and facilitates MiDAS to ensure complete genome duplication prior to cell division.

## RESULTS

### PCNA-K164 ubiquitination is required for DNA damage tolerance in human cells

Previous studies investigating the role of PCNA ubiquitination during DDT in human cells relied on knockdown of endogenous PCNA and overexpression of a K164R mutant or PCNA-ubiquitin fusion constructs (Niimi et al., 2008; Qin et al., 2013). To directly test the importance of this post-translational modification, we utilized CRISPR/Cas9 mediated gene targeting to knock-in an A>G mutation in the codon for K164 at the endogenous *PCNA* locus in RPE-1 cells, resulting in expression of a K164R mutant protein (Fig. S1 A). PCR analyses identified mono- and bi-allelic targeting of *PCNA* (Fig. S1, B and C), and subsequent Sanger sequencing confirmed one homozygous *PCNA^K164R^* (2B10) and two hemizygous *PCNA^K164R/-^* (A1 and B1) clones. Karyotype analysis revealed that *PCNA^K164R/-^* mutant cell lines had an inversion of one X chromosome at band q13q24 and a derivative chromosome X with the presence of chromosomal material of unknown origin at band q28 (Fig. S1 D). All mutant cell lines expressed similar steady-state levels of PCNA (Fig. S1 E) and equal amounts of chromatin-bound PCNA (Fig. 1 A). As expected, western blot analyses of PCNA ubiquitination after UV irradiation showed a dose-dependent increase in wildtype RPE-1 cells, but not in the *PCNA^K164R^* mutant cell lines (Fig. 1 A). Furthermore, we examined phosphorylated replication protein A (pRPA32) at S33 and phosphorylated histone H2AX (γH2AX) at S139 as a readout for replication stress and DNA double-stranded breaks (DSBs), respectively. We found a dose-dependent increase in pRPA32 and γH2AX in all cell lines with significantly elevated levels in the K164R mutants (Fig. 1 A). Due to the observed increases in replication stress and DSBs, we investigated the expression of DNA damage induced checkpoint markers phosphorylated p53 (S15) and p53. Although both markers showed a modest increase in the untreated mutant cell lines, p21 was not significantly activated (Fig. 1 B). These observations suggested that the lack of ubiquitination of PCNA at K164 confers an increased susceptibility to endogenous replication stress, but that endogenous DNA damage does not trigger a robust checkpoint response.

**Figure 1.**
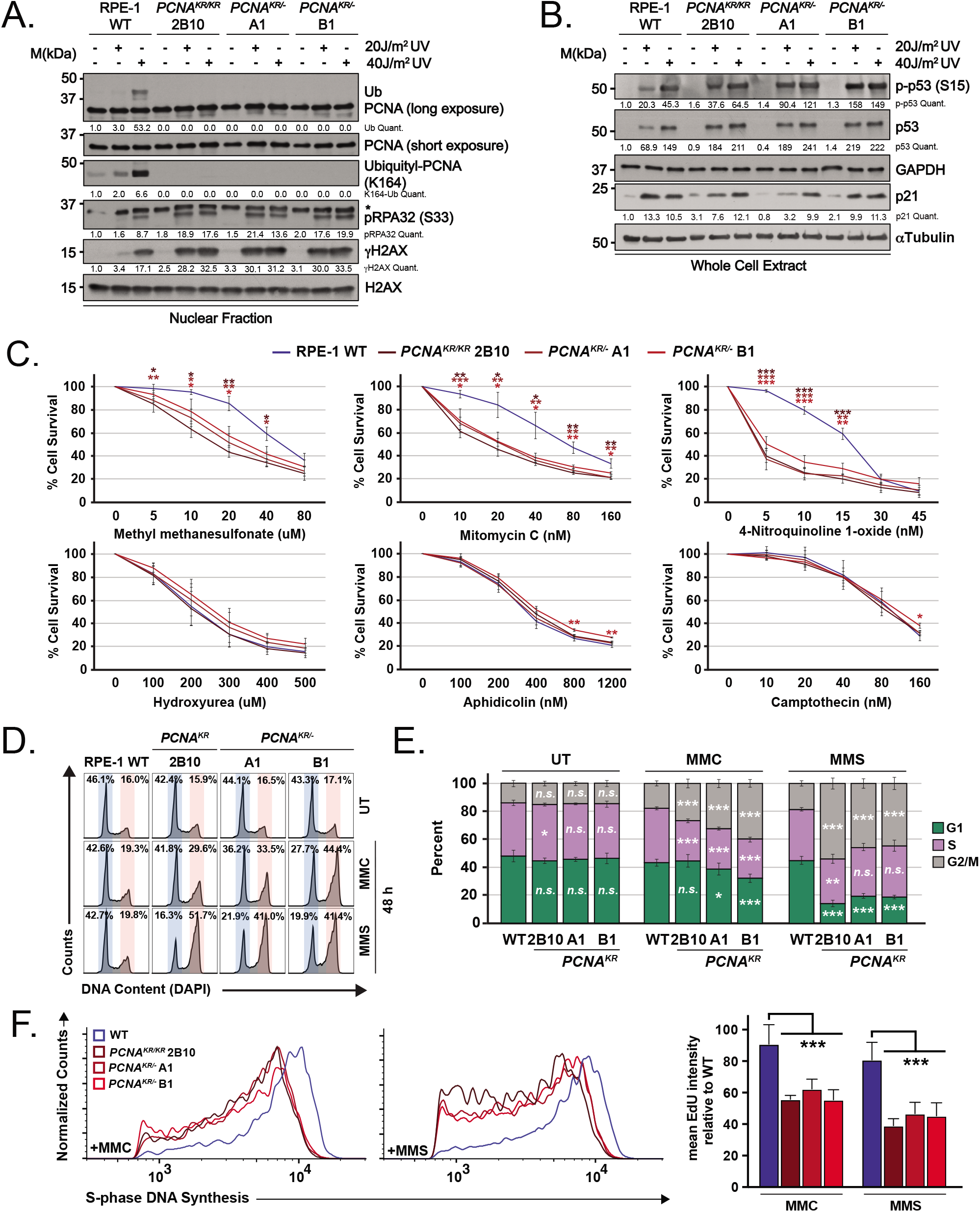
*PCNA^K164R^* mutant cell lines exhibit increased sensitivity to DNA damage. **A)** Chromatin associated PCNA, ubiquityl-PCNA (K164), phospho-RPA32 (S33) and γH2AX, with or without 20 J/m^2^ and 40 J/m^2^ UV treatment, with histone H2AX as the loading control. Band intensities were normalized to loading controls. **B)** Western blot analyses of whole cell extracts from wildtype RPE-1 and *PCNA^K164R^* cells for phospho-p53 (S15), p53, and p21 with or without 20 J/m^2^ and 40 J/m^2^ UV treatment, with GAPDH or tubulin as the loading control. Intensities of phospho-p53 (S15), p53 and p21 levels were normalized to loading controls. **C)** Comparison of drug sensitivity as measured by ATPase assay comparing average percentage viability in RPE-1 wildtype and *PCNA^K164R^* cell lines. Each drug and concentration tested is indicated. Error bars indicate standard deviation and statistical significance was calculated using students *t-test* with *>.05; **>.01, ***>.001; n = 9-12 replicate wells across three biological replicates for all data points. **D)** Representative cell cycle distribution of RPE-1 wildtype and *PCNA^K164R^* cell lines treated with MMC (20 nM) and MMS (20 µM) for 48 h, based on DNA content (DAPI). **E)** Cell cycle distribution of RPE-1 wildtype and *PCNA^K164R^* cell lines treated with MMC and MMS from three biological replicates. Percentage of each population in G1- (green), S- (purple) or G2/M-phase (gray) is shown. Error bars indicate standard deviation and statistical significance was calculated using students *t-test* with *>.05; **>.01, ***>.001; n = 6 replicate wells across three biological replicates. **F)** Histogram (left, middle) and quantification of mean fluorescent intensity (right) of EdU staining of S-phase cells from RPE-1 wildtype (blue) and *PCNA^K164R^* cells (maroon) treated with MMC and MMS; n = 6 across three biological replicates. Error bars indicate standard deviation and significance was calculated using two-way ANOVA with *>.05; **>.01, ***>.001.

To further understand the role of PCNA ubiquitination in DNA damage resistance, we utilized a luminescent (ATPase) cell viability assay to evaluate the sensitivity of *PCNA^K164R^* mutants to a variety of genotoxic drugs. The mutants were highly sensitive to methyl methanesulfonate (MMS), mitomycin C (MMC) and 4-nitroquinoline 1-oxide (4NQO), but not to hydroxyurea (HU), aphidicolin (APH) or camptothecin (CPT) (Fig. 1 C). To assess the effects of MMC and MMS on the cell cycle, we utilized quantitative chromatin flow cytometry. This allowed us to determine the cell cycle distribution using DAPI-staining for total DNA content in combination with a 30 minute 5-ethynyl-2’-deoxyuridine (EdU) pulse to label S phase cells. Cell cycle analysis revealed a G2 arrest in the mutants when challenged with MMC or MMS (Fig. 1, D and E) and a significant decrease in S phase DNA synthesis (Fig. 1 F). Taken together, these data indicate that PCNA-K164 ubiquitination is required in RPE-1 cells for tolerance of lesion-induced replication stress.

### *PCNA^K164R^* mutants exhibit decreased global DNA synthesis caused by impaired origin activation

Although DAPI-staining of genomic DNA did not reveal significant cell cycle profile changes between RPE-1 wildtype and *PCNA^K164R^* mutants (Fig. 1, D and E), the mutant cell lines exhibited a significant decrease in cell proliferation (Fig. 2 A) that could not be explained by the modest increase we measured in apoptosis by flow cytometry of cells stained with propidium iodide (PI) and annexin V (Fig. S1 F). A similar proliferation defect was observed in *PCNA^K164R^* mutant 293T cell lines (Fig. S2 A) (Thakar et al., 2020), whereas this has not been reported in budding yeast, chicken, or mouse cell lines (Hoege et al., 2002; Simpson et al., 2006; Arakawa et al., 2006; Langerak et al., 2007). Thus, we more carefully assessed whether DNA replication was slower in the K164R mutants using quantitative chromatin flow cytometry (Fig. 2 B; and Fig. S2 B). We observed small changes in the cell cycle distribution of RPE-1 *PCNA^K164R^* cells compared to wildtype (Fig. 2 C), associated with a mild but significant defect in EdU incorporation in two of the three mutant clones (Fig. 2 D). The increase in the G2 population, usually interpreted as a defect in completing DNA replication, was proportional to the decrease in G1 phase (Fig. 2, B and C). Both the increase in G2 population and defective DNA synthesis were more pronounced in 293T *PCNA^K164R^* mutants (Fig. S2, B-D). The 293T *PCNA^K164R^* mutant complemented with wildtype PCNA, however, did not fully rescue these phenotypes. We suspected that this was due to the presence of PCNA trimers composed entirely or partially of K164R mutant PCNA, which renders these trimers non- or only partially functional for DDT.

**Figure 2.**
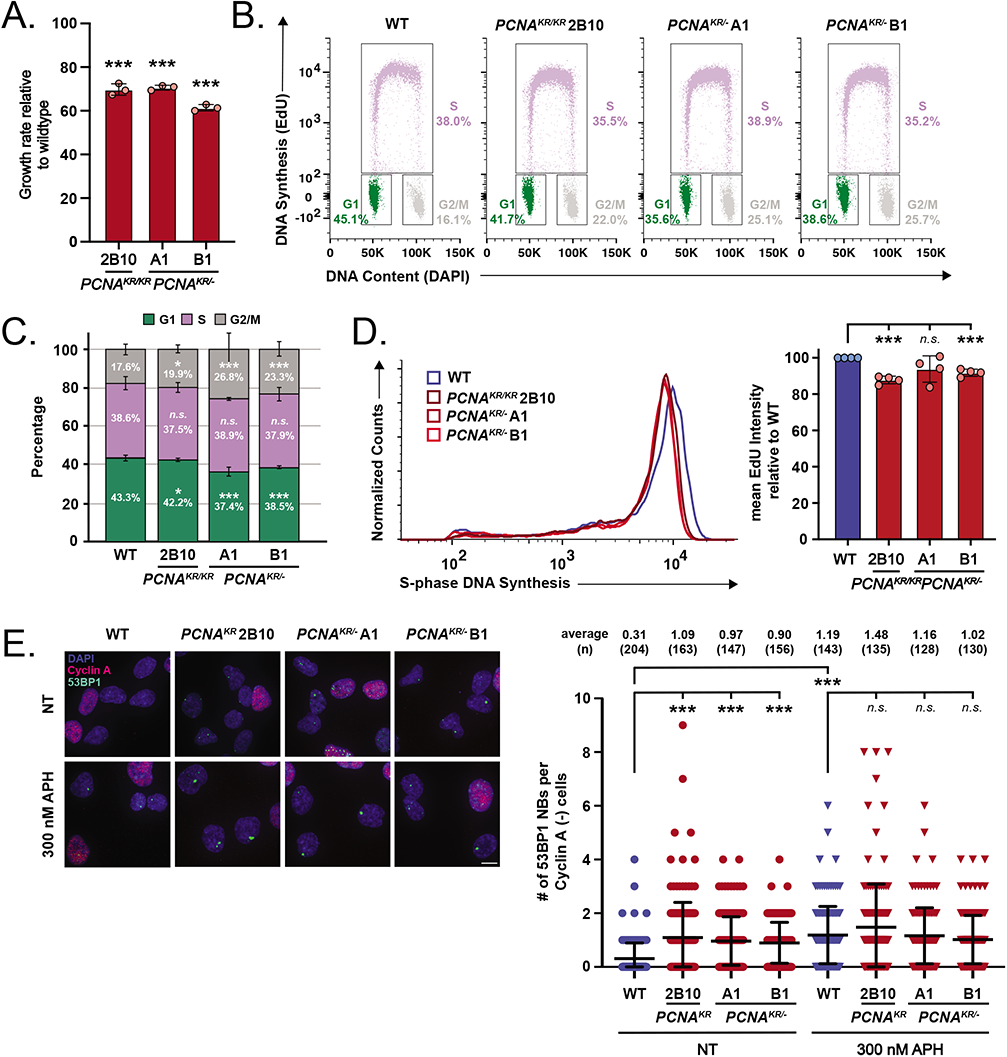
The PCNA-K164R mutation causes DNA synthesis defects during unperturbed replication. **A)** Average cell proliferation rate in *PCNA^K164R^* cell lines normalized to wildtype. For each cell line n = 9 wells across three biological replicates. Error bars indicate standard deviation and significance was calculated using one-way ANOVA with *>.05; **>.01, ***>.001. **B)** Representative cell cycle distribution of RPE-1 wildtype and *PCNA^K164R^* cell lines based on DNA content (DAPI) and DNA synthesis (EdU incorporation). Percentage of each population in G1- (green), S- (purple) or G2/M-phase (gray) is shown. **C)** Cell cycle distribution of RPE-1 wildtype and *PCNA^K164R^* cell lines from four biological replicates. Percentage of each population in G1- (green), S- (purple) or G2/M-phase (gray) is shown. Error bars indicate standard deviation and significance was calculated using students *t-test* with *>.05; **>.01, ***>.001. **D)** Histogram (left) and quantification of mean fluorescent intensity (right) of EdU staining of S-phase cells from RPE-1 wildtype (blue) and *PCNA^K164R^* cells (maroon); n=12 across four biological replicates. Error bars indicate standard deviation and significance was calculated using students *t-test* with *>.05; **>.01, ***>.001. **E)** (Left) Image of 53BP1 NB and cyclin A staining in RPE-1 cell lines. DAPI (blue), 53BP1 NB (green), and cyclin A (pink) are indicated. Scale bar at 20 µm. (Bottom) 53BP1 NB quantification of untreated (circles) and APH treated (triangles) in wildtype (blue) and *PCNA^K164R^* lines (maroon). Number (n) of nuclei quantified is listed. Error bars indicate standard deviation and significance was calculated using Kruskal-Wallis with Dunn’s multiple comparison test.

We reasoned that defects in DNA replication might result in a high frequency of under-replicated DNA that could persist into mitosis. One fate of under-replicated DNA is the conversion into DNA lesions that are sequestered into large chromatin domains known as 53BP1 NBs and inherited by daughter cells (Harrigan et al., 2011; Lukas et al., 2011; Moreno et al., 2016; Lezaja and Altmeyer, 2018). These lesions are repaired in the following S phase (Spies et al., 2019). To investigate whether *PCNA^K164R^* mutants have elevated levels of under-replicated DNA, we analyzed 53BP1 NB formation in G1 nuclei. *PCNA^K164R^* mutants had a 3-fold increase in 53BP1 NBs under unperturbed conditions (Fig. 2 E; and Fig. S2 E). Low-level replication stress induced by APH treatment led to a 3-to-5-fold increase in 53BP1 NB formation in the wildtype cells; however, there was no additional increase in NBs in the mutant cells. Given that *PCNA^K164R^* mutants have a similar sensitivity to APH as the wildtype cells (Fig. 1 C), the lack of an increase in 53BP1 NB formation suggested that mutant cells: (1) experience maximal replication stress under steady-state conditions, and/or (2) resolve LRIs via alternative mechanism(s). Interestingly, the 293T *PCNA^K164R^* mutant complemented with wildtype PCNA partially rescued APH induced 53BP1 NBs (Fig. S2 E). These results reveal that loss of K164 ubiquitination negatively affects DNA synthesis under unperturbed conditions, especially in 293T cells. These results are consistent with our previous report, suggesting that K164 ubiquitination has a role during normal DNA replication in human cells (Thakar et al., 2020),

We also measured chromatin-bound MCM2, as a marker for origin licensing in G1 phase (Fig. 3 A) (Forment and Jackson, 2015; Matson et al., 2017). Surprisingly, we observed a 15-20% reduction in MCM2 loading in the RPE-1 *PCNA^K164R^* mutants when we quantified the percentage of MCM2-positive cells in late G1 phase just prior to S phase entry (Fig. 3, B and D). This decrease was not due to an overall reduction in steady-state levels of MCM2 (Fig. S1 E). To quantify chromatin-bound MCM2 in early S phase in more detail, we focused on the populations within the black rectangles shown in Fig. 3C. As described previously, we defined the cutoff for fully licensed cells in wildtype and applied the same cutoff to the mutant samples (Matson et al., 2019). Cells in early S phase that fell below that cutoff were defined as under-licensed (Fig. 3 C, dashed line). These populations in the *PCNA^K164R^* mutants had a broader range of chromatin-bound MCM2 with a 30% increase in under-licensed cells when compared to wildtype (Fig. 3 E). A similar defect was observed in the 293T *PCNA^K164R^* mutant (Fig. S3, A and B). Together, these data suggest a previously unknown connection between the PCNA-K164R mutation and origin licensing.

**Figure 3.**
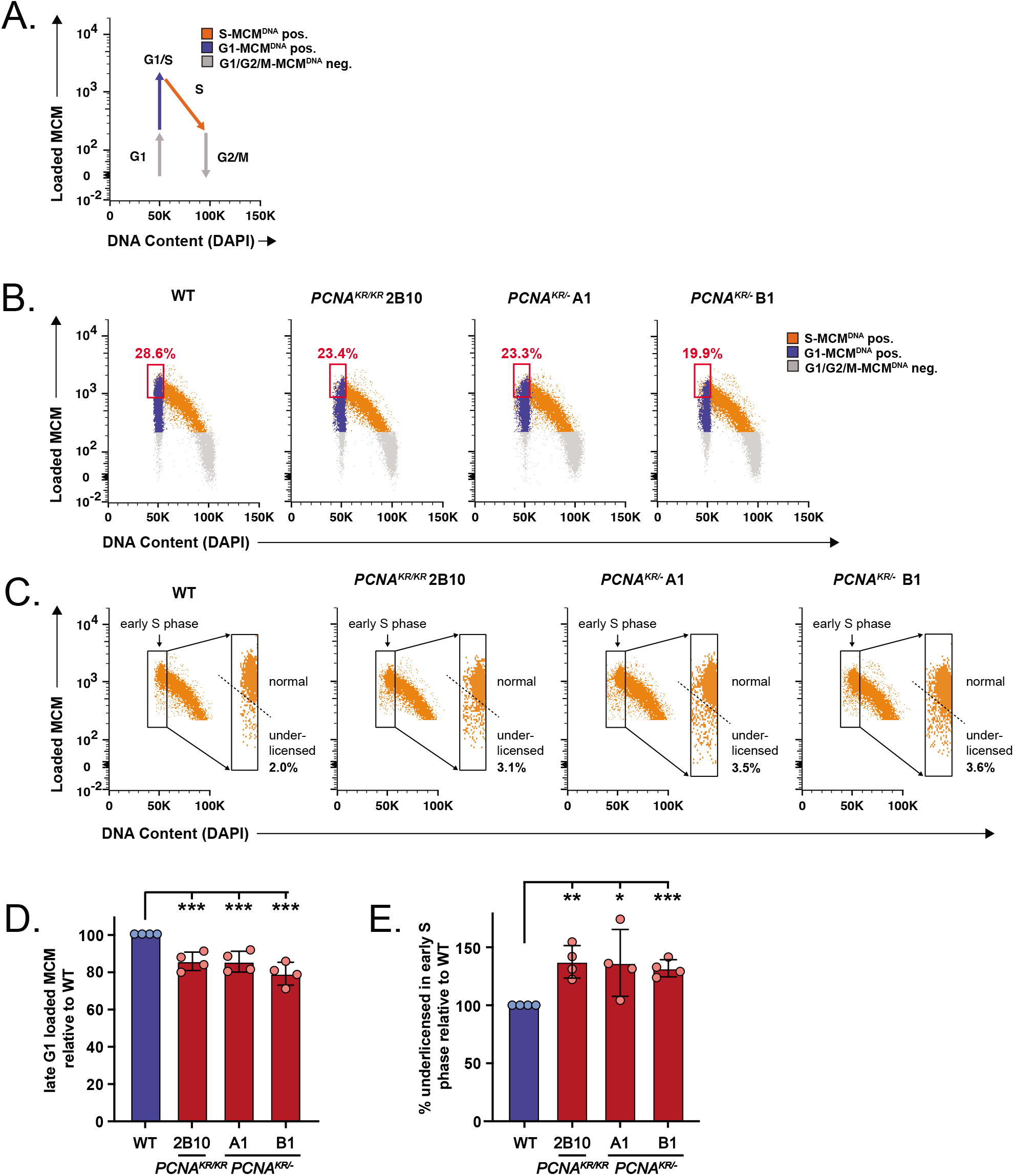
Defects in DNA synthesis in RPE-1 *PCNA^K164R^* cells impinges on origin licensing. **A)** Schematic of quantitative chromatin flow cytometry analysis. Cell cycle phase is defined by DNA content, EdU incorporation and chromatin loaded MCM2. **B)** Representative chromatin flow cytometry plots for RPE-1 wildtype and *PCNA^K164R^* cells. G1-phase/MCM positive cells (blue), S-phase/MCM positive cells (orange) and G1- or G2/M-phase/MCM negative cells (gray) are indicated. Percentage of MCM2 stained cells in late G1 is indicated (red box). **C)** Gating of early S cells from B, showing only the S phase, MCM/DNA-positive cells. Rectangles define early S phase cells as MCM/DNA-positive with 2C DNA content. The dashed line defines the cutoff between normally licensed versus under-licensed cells. **D)** Quantification of the percentage of MCM2 stained cells in late G1 from RPE-1 wildtype (blue) and *PCNA^K164R^* cells (maroon); n=12 across four biological replicates. Error bars indicate standard deviation and significance was calculated using students *t-test* with ***>.001. **E)** Percentage of under-licensed cells in early S phase from RPE-1 wildtype (blue) and *PCNA^K164R^* cells (maroon); n = 12 across four biological replicates. Error bars indicate standard deviation and significance was calculated using students *t-test* with *>.05, **>.01, ***>.001.

### PCNA-K164R mutants exhibit a decrease in the number of active replication forks

To better understand the DNA synthesis defects in *PCNA^K164R^* mutants, we performed DNA combing analysis under unperturbed conditions. We sequentially pulse-labeled wildtype and *PCNA^K164R^* cells with a thymidine analog, IdU (green) for 20 minutes followed by a second thymidine analog, CldU (red) for 20 minutes. Due to reduced EdU incorporation and origin licensing in the mutants, we predicted that fewer replication origins would translate into longer inter-origin distances (IODs). The average IOD in wildtype cells was ∼102 kb whereas the average in *PCNA^K164R^* RPE-1 mutants were markedly increased to ∼136-145 kb (Fig. 4, A and B). This difference equates to ∼33-42% fewer active origins. In line with these data, we measured a 20-30% decrease in new origin firing events in all *PCNA^K164R^* mutants (Fig. 4, A and C; and Fig. S3 C). Although the mutants displayed a significantly lower number of replication forks, fork stability was not compromised (Fig. 4 D) and only one mutant exhibited spontaneous fork stalling (Fig. 4 E). These findings reveal that incomplete DNA replication as indicated by 53BP1 NBs is likely caused by diminished origin activation, which translates into fewer active replication forks.

**Figure 4.**
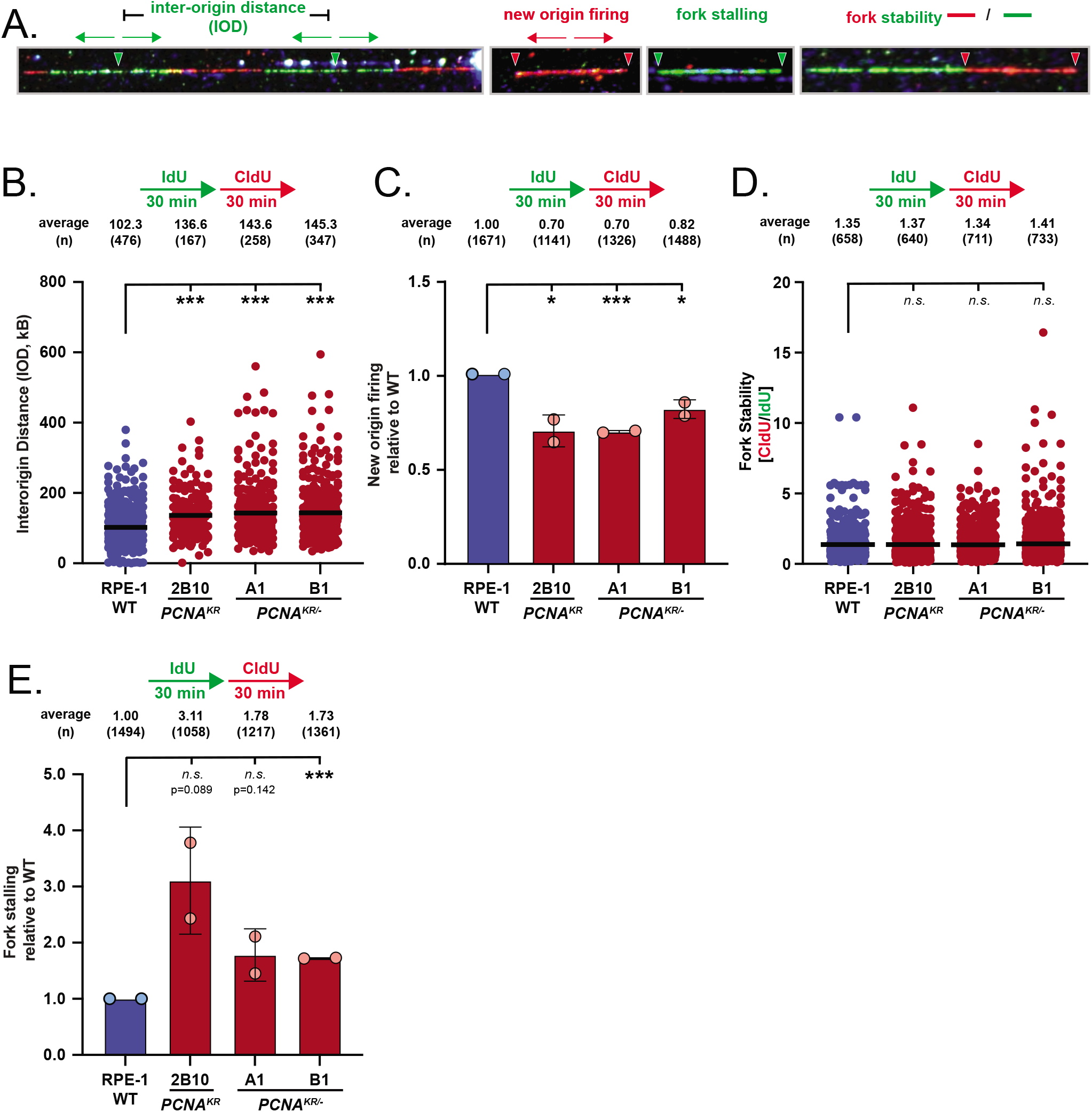
PCNA-K164R decreases the number of active replication forks. **A)** Example of fibers used for DNA combing analyses. Active replication forks were sequentially labeled with IdU (25 µM, green) for 30 minutes followed by labeling with CldU (200 µM, red) for 30 minutes. Inter-origin distance (IOD) is measured as center-to-center distance between two adjacent progressing bidirectional forks. Green arrows represent the direction of fork progression. New origin firing (NOF) is measured as CldU-only tracts divided by the total number of tracts. Fork stalling is measured as IdU-only tracts divided by the total number of tracts. Fork stability is calculated by dividing the CldU tract length by the IdU tract length. All quantifications (B-E) were obtained under unperturbed conditions. **B)** IOD quantification from two biological replicates in RPE-1 wildtype (blue) and *PCNA^K164R^* cells (maroon). Average IOD and number (n) quantified is listed. Significance was calculated by Kruskal-Wallis with Dunn’s multiple comparison test with ***<.001. **C)** New origin firing (NOF) events from two biological replicates in RPE-1 wildtype (blue) and *PCNA^K164R^* cells (maroon). Number (n) of events quantified is listed. Significance was calculated using students *t-test* with *>.05, **>0.01, ***>0.001. **D)** Fork stability from two biological replicates in RPE-1 wildtype (blue) and *PCNA^K164R^* cells (maroon). Average fork stability and number (n) quantified is listed. Significance was calculated by Kruskal-Wallis with Dunn’s multiple comparison test. **E)** Fork stalling events from two biological replicates in RPE-1 wildtype (blue) and *PCNA^K164R^* cells (maroon). Number (n) of events quantified is listed. Significance was calculated using students *t-test* with ***>0.001.

### PCNA-K164 is an important regulator of FANCD2 mono-ubiquitination and MiDAS

Under-replicated genomic regions can increase LRIs, leading to the formation of FANCD2 foci during G2/M phase of the cell cycle and activating MiDAS as the cells’ final attempt to complete DNA replication prior to cell division (Graber-Feesl et al., 2019). To visualize MiDAS, we pulse-labeled cells with EdU and examined its incorporation and co-localization with FANCD2 in prophase/prometaphase nuclei (Bhowmick et al., 2016; Graber-Feesl et al., 2019). Knockdown of MCM4, an essential component of the replicative helicase, decreases origin licensing genome-wide (Ge et al., 2007; Ibarra et al., 2008; Alver et al., 2014) and led to increased EdU and FANCD2 foci (Fig. S4 A-C), consistent with previous data following the depletion of origin recognition complex subunit 1 (Graber-Feesl et al., 2019). Given the altered replication dynamics in *PCNA^K164R^* cells (Fig. 2 and 4), we suspected that these mutants would upregulate MiDAS to minimize under-replicated DNA passing through mitosis. Surprisingly, RPE-1 *PCNA^K164R^* cells had significantly fewer FANCD2 foci (∼2-3-fold) and a corresponding decrease in MiDAS (∼1.5-3.5-fold) compared to wildtype cells (Fig. 5 A-C; and Fig. S5 B). Similar results were seen in the 293T *PCNA^K164R^* mutant. Interestingly, the 293T *PCNA^K164R^* mutant complemented with wildtype PCNA partially restored FANCD2 foci formation, however, it was not sufficient to fully rescue the MiDAS defect (Fig. 5 D and E; and Fig. S5 C). These findings suggest a role for PCNA-K164 ubiquitination in regulating MiDAS through the recruitment and/or mono-ubiquitination of FANCD2.

**Figure 5.**
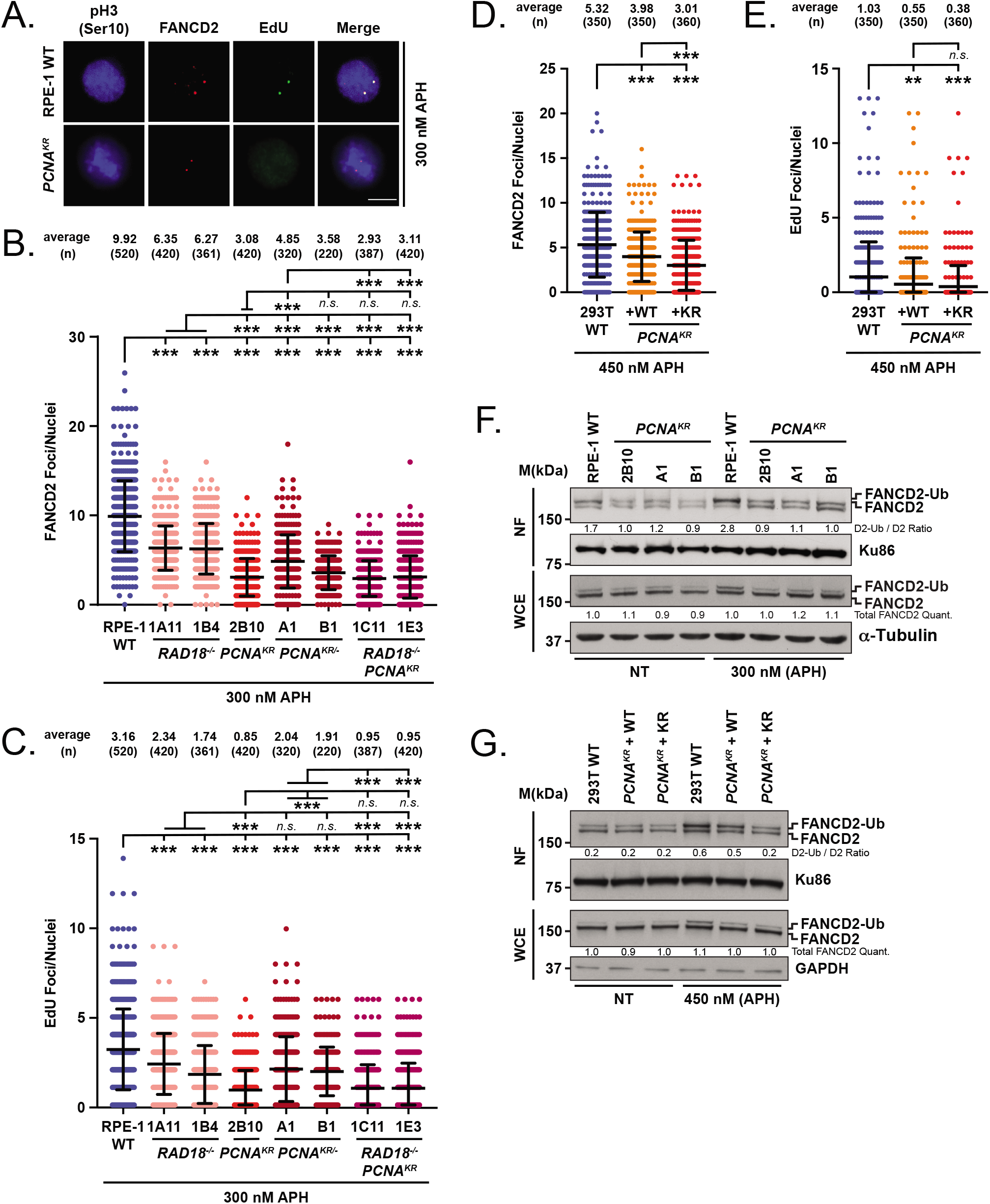
PCNA-K164 regulates FANCD2 mono-ubiquitination and MiDAS. **A)** Representative images of phospho-H3-stained nuclei/chromosomes (blue), EdU foci (green), and FANCD2 (red) foci for each RPE-1 cell line. Scale bars are 5 µm. **B)** FANCD2 foci quantification from two biological replicates in RPE-1 wildtype (blue), *RAD18^-/-^* (pink), *PCNA^K164R^* (red), *PCNA^K164R/-^* (maroon) and *RAD18^-/-^PCNA^K164R^* (purple) cells treated with APH. Number (n) of nuclei quantified is listed. Significance was calculated by Kruskal-Wallis with Dunn’s multiple comparison test with ***<.001. **C)** EdU foci quantification from two biological replicates in RPE-1 wildtype (blue), *RAD18^-/-^* (pink), *PCNA^K164R^* (red), *PCNA^K164R/-^* (maroon) and *RAD18^-/-^PCNA^K164R^* (purple) cells treated with APH. Number (n) of nuclei quantified is listed. Significance was calculated by Kruskal-Wallis with Dunn’s multiple comparison test with ***<.001. **D)** FANCD2 foci quantification from two biological replicates in 293T wildtype (blue) and *PCNA^K164R^* cells complemented with either wildtype (orange) or a K164R (red) cDNA treated with APH. Number (n) of nuclei quantified is listed. Significance was calculated by one-way ANOVA with Tukey’s multiple comparison test with ***<.001. **E)** EdU foci quantification from two biological replicates in 293T wildtype (blue) and *PCNA^K164R^* cells complemented with either wildtype (orange) or a K164R (red) cDNA treated with APH. Number (n) of nuclei quantified is listed. Significance was calculated by one-way ANOVA with Tukey’s multiple comparison test with ***<.001. **F)** (Top) Chromatin associated FANCD2 with Ku86 as the loading control in RPE-1 wildtype and *PCNA^K164R^* cell lines. Ratio of mono-ubiquitinated to non-ubiquitinated FANCD2 is indicated. (Bottom) Western blot analyses of whole cell extracts from RPE-1 wildtype and *PCNA^K164R^* cell lines for FANCD2 with α-tubulin as the loading control. Intensities of total FANCD2 protein was normalized to loading controls. **G)** (Top) Chromatin associated FANCD2 with Ku86 as the loading control in 293T wildtype and *PCNA^K164R^* cells complemented with either wildtype or a K164R cDNA. Ratio of mono-ubiquitinated to non-ubiquitinated FANCD2 is indicated. (Bottom) Western blot analyses of whole cell extracts from 293T wildtype and *PCNA^K164R^* cells complemented with either wildtype or a K164R cDNA for FANCD2 with GAPDH as the loading control. Intensities of total FANCD2 protein was normalized to loading controls.

We previously showed that depletion of FANCA, a component of the FA core complex involved in the mono-ubiquitination of FANCD2, significantly impaired MiDAS (Graber-Feesl et al., 2019). Furthermore, since mono-ubiquitinated FANCD2 binds tightly to DNA (Alcón et al., 2020; Wang et al., 2020) and is a prerequisite for focus formation, we examined whether K164 ubiquitination correlated with FANCD2 mono-ubiquitination. Under unperturbed conditions, FANCD2 mono-ubiquitination was drastically reduced in the RPE-1 *PCNA^K164R^* cells. Interestingly, upon inducing low-level replication stress by exposure to APH, FANCD2 mono-ubiquitination was elevated in the RPE-1 and 293T wildtype cells, but not in the *PCNA^K164R^* mutants (Fig. 5 F and G, NF). The decrease in FANCD2 mono-ubiquitination observed in these mutants was not due to an overall reduction in steady-state levels of FANCD2 (Fig. 5 F and G, WCE).

RAD18-mediated PCNA-K164 ubiquitination has been linked to FANCD2 mono-ubiquitination (Geng et al., 2010; Song et al., 2010). Thus, we generated *RAD18^-/-^* single and *RAD18^-/-^:PCNA^K164R^* double mutants (Fig. S5 A) and examined them for mitotic EdU incorporation and FANCD2 foci. *RAD18^-/-^* cells displayed approximately 40% fewer EdU and FANCD2 foci compared to wildtype. However, *RAD18^-/-:^PCNA^K164R^* double mutants had similar levels of EdU and FANCD2 foci as the *PCNA^K164R^* single mutant, consistent with RAD18 acting upstream of PCNA ubiquitination (Fig. 5 A-C; and Fig. S5 B). The number of FANCD2 foci in *RAD18^-/-^* cells was slightly elevated compared to that in *PCNA^K164R^* and *RAD18^-/-:^PCNA^K164R^* double mutants (Fig. 5 B). These results are in agreement with reports that three additional E3 ubiquitin ligases, ring finger protein 8 (RNF8), Cullin-4-RING-ligase (CRL4)-Ddb1-Cdt2 (CRL4^Cdt2^) and ring finger and WD repeat domain 3 (RFWD3) are capable of mono-ubiquitinating PCNA (Zhang et al., 2008; Terai et al., 2010; Gallina et al., 2021). Therefore, in the absence of RAD18, PCNA ubiquitination by alternative E3 ligases can partially activate MiDAS. In contrast, when PCNA is unable to be ubiquitinated at K164, MiDAS is severely compromised. Taken together, these data indicate that PCNA-K164 plays a critical role in activating MiDAS in response to under-replicated DNA through modulating mono-ubiquitination of FANCD2.

### Defective MiDAS in *PCNA^K164R^* cells results in anaphase abnormalities

When MiDAS fails, the persistence of LRIs in mitosis results in the formation of chromatin bridges during anaphase (Chan and Hickson, 2011; Chan et al., 2018). If these bridges persist into late anaphase they can: (1) cause perturbations to the spindle apparatus, facilitating the formation of lagging chromosomes (Chan et al., 2018), (2) be physically sheared by the force of the spindle apparatus, resulting in DSBs, aberrant chromosome segregation and possibly chromothripsis, and/or (3) prevent cytokinesis, thus leading to bi-nucleated cells (Crasta et al., 2012; Hayashi and Karlseder, 2013; Hatch and Hetzer, 2015). To investigate whether *PCNA^K164R^* mutants harbor elevated levels of under-replicated DNA during mitosis, we analyzed the frequency of anaphase abnormalities present as DAPI-positive chromatin bridges and lagging chromosomes. Low-level replication stress induced by APH treatment led to a significant increase in the percentage of abnormal anaphases (∼40%) in the *PCNA^K164R^* mutants (Fig. 6 A and B), exacerbating the replication defect of the mutants. The finding that the mutants are not more sensitive to APH than wildtype cells (Fig. 2C) further suggests that they tolerate these chromosome structures, likely at the expense of genome alterations, such that overall survival is not diminished.

**Figure 6.**
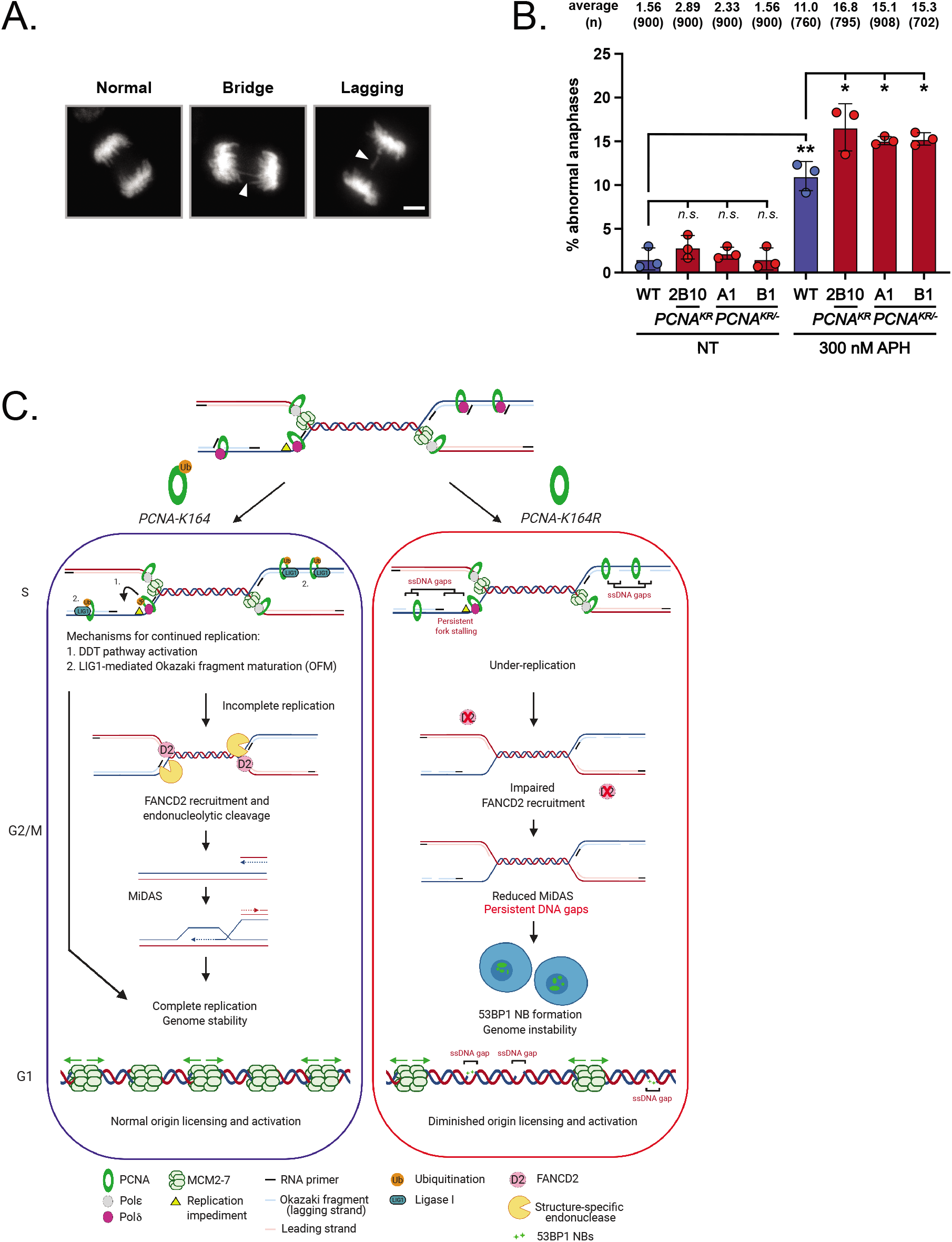
Defective MiDAS in *PCNA^K164R^* cells results in anaphase abnormalities. **A)** Representative images of DAPI stained chromosomes during anaphase: (left) normal, (middle) chromatid bridge, and (right) lagging chromosome. Scale bars are 5 µm. **B)** Percentage of abnormal anaphases in untreated (lanes 1-4) and APH treated (lanes 5-6) in wildtype (blue) and *PCNA^K164R^* lines (maroon). Number (n) of anaphases quantified is listed. Error bars indicate standard deviation and significance was calculated using students *t-test* with *>.05; **>.01, ***>.001. **C)** Wildtype (Left): PCNA K164-ubiquitination occurs during normal S phase to promote gap filling during OF maturation. If the replication fork encounters a DNA lesion/impediment, cells can complete replication by activating DDT pathways. DNA that is not duplicated before cells enter G2/M phase stimulates MiDAS through the recruitment of FANCD2. In the subsequent G1 phase, origins are licensed in G1 and activated in S-phase. *PCNA^K164R^* (Right): During S phase, in the absence of PCNA-K164 ubiquitination, OF maturation is impaired, which leads to the accumulation of ssDNA gaps. In the presence of a DNA lesion/impediment, DDT pathway activation does not occur, and under-replicated DNA accumulates (ssDNA gaps). These under-replicated regions will persist into mitosis, leading to anaphase abnormalities because MiDAS is impaired. In the subsequent G1 phase, ssDNA gaps interfere with origin licensing, resulting in diminished origin activation.

## DISCUSSION

Here, we describe novel roles for PCNA-K164 modification in maintaining genome stability by resolving LRIs through MiDAS and promoting efficient origin activation (Fig. 6 C). We recently provided evidence that in the absence of PCNA ubiquitination during unperturbed conditions, ssDNA gaps accumulate during lagging strand synthesis and cause defects in OF maturation (Thakar et al., 2020). We propose that these gaps interfere with DNA replication in subsequent cell cycles (Fig. 2 D; and Fig. S2 D). In this study, we corroborate that under-replicated DNA persists into the next cell cycle (Fig. 2 E; and Fig S3 E) due to severely compromised MiDAS (Fig. 5 B-E) because of reduced FANCD2 mono-ubiquitination (Fig. 5 F and G). Furthermore, the increase in ssDNA gaps from previous rounds of DNA replication caused insufficient origin licensing in G1 phase (Fig. 3; and Fig. S3 A and B), thereby perpetually exacerbating the under-replication phenotype.

### PCNA K164 ubiquitination prevents the generation of ssDNA gaps by multiple mechanisms

Our studies demonstrate that the inability to ubiquitinate PCNA at K164 severely inhibited faithful genome duplication. Consistent with previous studies in mammalian cells, *PCNA^K164R^* mutants were sensitive to DNA lesions induced by UV, MMS, and MMC (Fig. 1) (Niimi et al., 2008; Hendel et al., 2011; Qin et al., 2013; Vujanovic et al., 2017). However, we did not detect additional sensitivity to CPT. This finding is similar to an earlier study in human cells (Niimi et al., 2008), but is contrary to studies of *PCNA^K164R^* mouse embryonic fibroblasts (Vujanovic et al., 2017). This discrepancy could be due to organismal differences. A K164R mutation has never been identified in human tissues, but homozygous *PCNA^K164R^* mice were viable (Langerak et al., 2007). However, these mice were sterile, exhibited bone marrow (BM) failure and defects in somatic hypermutation and class switch recombination (Langerak et al., 2007; Roa et al., 2008; Pilzecker et al., 2017), implying that K164 ubiquitination is not completely dispensable in mice. Interestingly, infertility and BM failure are phenotypes shared between *PCNA^K164R^* and FA mouse models (Parmar et al., 2009; Bakker et al., 2013), consistent with the functional redundancy of the respective pathways in suppressing ssDNA gaps during lagging strand synthesis (Thakar et al., 2020).

The sensitivity of *PCNA^K164R^* mutant cells to lesion-induced DNA damage (Fig. 1) (Thakar et al., 2020) suggests that these cells are defective for TLS. Multiple groups have demonstrated that TLS occurs on both the leading and lagging strands (Pagès et al., 2008; Yoon et al., 2009, 2012; Hedglin and Benkovic, 2017), where its primary function is to suppress ssDNA gap formation (Nayak et al., 2020). In the absence of TLS polymerase η, MiDAS increases significantly (Garcia-Exposito et al., 2016; Bergoglio et al., 2013). Elevated MiDAS could be a compensatory mechanism to counteract the increase in ssDNA gaps. Our data suggests that ubiquitination of PCNA at K164 prevents the accumulation of ssDNA regions by promoting both TLS and MiDAS (Fig. 5 B-E).

Several lines of evidence support that PCNA ubiquitination at K164 prevents ssDNA gap formation by promoting OF maturation. In budding yeast, loss of *CDC9* (LIG1), *RAD27* (FEN1), and the PCNA unloader *ELG1* (ATAD5), trigger PCNA ubiquitination (Das-Bradoo et al., 2010; Nguyen et al., 2013; Becker et al. 2015). A synthetic genetic array screen using PCNA-K164R mutants as queries revealed a genetic interaction signature similar to that of strains defective in lagging strand synthesis including *RFC5*, *POL31* (Pol δ), *RAD27*, and *ELG1* mutants (Becker et al., 2015). Human *PCNA^K164R^* cells phenocopy the DNA synthesis defects seen in LIG1 and FEN1 depleted cells, further supporting a conserved role for PCNA ubiquitination in promoting OF processing by gap suppression (Thakar et al., 2020). Taken together, our data further substantiate the notion that PCNA-K164 ubiquitination promotes complete genome duplication by preventing ssDNA accumulation caused by defective TLS and/or OF maturation.

### Helicase loading and origin activation are impaired in PCNA K164R mutants

There is a growing body of evidence that suggests that PCNA-K164 ubiquitination is not only induced by DNA damage, but occurs during normal DNA replication (Frampton et al., 2006; Leach and Michael, 2005; Simpson et al., 2006; Thakar et al., 2020). In the absence of exogenous replication stress, we show that both RPE-1 and 293T *PCNA^K164R^* mutants have a significant reduction in proliferation rate (Fig. 2 A; and Fig. S2 A), and a subtle, but significant global DNA synthesis defect (Fig. 2D; and Fig. S2 D). Unexpectedly, the primary replication defects in *PCNA^K164R^* cells were reduced origin licensing and origin firing (Fig. 3 E; Fig. 4 C; and Fig. S3 B and C). In agreement with these observations, we show that *PCNA^K164R^* cells have increased IODs (Fig. 4 B).

Origin licensing studies in yeast revealed that MCM2-7 hexamers are loaded cooperatively in a head-to-head orientation onto double-stranded (ds) DNA (Remus et al., 2009; Evrin et al., 2009; Ticau et al., 2015). Origin activation in budding yeast requires MCM10 (Ricke and Bielinsky, 2004), which promotes dsDNA unwinding and allows the head-to-head CMG complexes to bypass one another on ssDNA (Langston and O’Donnell, 2019). Thus, replication initiation only occurs at origins where the replicative helicase is appropriately loaded onto dsDNA. In *PCNA^K164R^* mutants, the accumulation of ssDNA might impair origin licensing (Fig. 6C). Furthermore, the defects in OF processing in *PCNA^K164R^* cells leads to the accumulation of chromatin-bound PAR chains (Thakar et al., 2020), which are added by poly(ADP-ribose) polymerase (PARP), a sensor of unligated OF fragments (Hanzlikova et al., 2018). These chains might provide an additional barrier to robust MCM2-7 loading. Considering the MCM2 loading defect (Fig. 3; and Fig. S3 A and B) and increased under-replication (Fig. 2 E) observed in the *PCNA^K164R^* cells, we propose that accumulation of ssDNA gaps prevents efficient loading of MCM2-7 double hexamers in the subsequent G1 phase (Fig. 6 C). Future studies utilizing genome-wide ligation of 3’-OH ends followed by sequencing (GLOE-Seq) would allow for mapping and quantification of these gaps to delineate whether specific genomic loci are affected in *PCNA^K164R^* mutants (Sriramachandran et al., 2020).

G1 phase 53BP1 NBs delay replication of under-replicated loci until late S phase by recruiting replication timing regulatory factor 1 (RIF1) (Spies et al., 2019). This process protects these regions from aberrant processing and genotoxic RAD51-mediated recombination. During late S phase, RAD52 mediates the complete duplication of under-replicated loci. Thus, an alternative but not mutually exclusive model is that in the absence of PCNA-K164 ubiquitination, under-replicated DNA that is not resolved by MiDAS is packaged into 53BP1 NBs, which directly inhibit origin licensing.

### MiDAS activation by FANCD2 is dependent on PCNA ubiquitination

Our observation that *PCNA^K164R^* mutants are defective in MiDAS reveals an unexpected role for K164 ubiquitination (Fig. 5 B-E, and Fig. S5 B and C). A similar defect was observed in *RAD18^-/-^* cells (Fig. 5 B and C), whereas *RAD18^-/-^:PCNA^K164R^* double mutants had no additional defect, confirming that RAD18 and PCNA function in the same pathway. Furthermore, RAD18-dependent K164 ubiquitination accounts for only ∼40% of the MiDAS defect observed in *PCNA^K164R^* cells, implying that other E3 ligases modify K164 to activate MiDAS. Future studies will be needed to address whether RNF8, CRL4^Cdt2^, RFWD3 or an unknown E3 ligase functions in this capacity.

FANCD2 is a key regulator of MiDAS in noncancerous cells (Graber-Feesl et al., 2019). We demonstrate here that APH treated *PCNA^K164R^* cells exhibit reduced levels of chromatin bound mono-ubiquitinated FANCD2, although total FANCD2 levels were unaltered (Fig. 5 F and G). Based on these observations, we speculate that FANCD2 is recruited to under-replicated regions by PCNA. Consistent with our data, PCNA and FANCD2 co-localize following HU-and APH-induced replication stress (Hussain et al., 2004; Howlett et al., 2005). One possibility is that PCNA directly recruits FANCD2 through its PCNA-interacting peptide (PIP) box (Howlett et al., 2009), and/or through its coupling of ubiquitin conjugation to endoplasmic reticulum degradation (CUE) domain (Rego et al., 2012). Recent studies demonstrated that FANCD2 exists as a homodimer, and in response to replication stress one subunit is exchanged for FANCI, forming a FANCD2-FANCI (D2-I) heterodimer. Ubiquitination of DNA bound D2-I complex by the FA core complex locks it onto DNA (Alcón et al., 2020; Wang et al., 2020). Specifically, FANCD2 is mono-ubiquitinated by FANCL (Meetei et al., 2003; Alpi et al., 2008). Limited evidence suggests that FANCL recruitment may require RAD18-mediated PCNA-K164 ubiquitination, which in turn stimulates FANCD2 mono-ubiquitination (Geng et al., 2010). Future studies are required to test whether PCNA-K164 ubiquitination mediated recruitment of FANCL is critical for the mono-ubiquitination of FANCD2 and required for its stable association with under-replicated loci.

## MATERIALS AND METHODS

### Cell lines

RPE-1 cells were grown in Dulbecco’s Modified Eagle Medium: Nutrient Mixture F12 (DMEM/F12, Gibco 11320) supplemented with 10% fetal bovine serum (FBS, Sigma F4135) and 1% Penicillin-Streptomycin (Pen Strep, Gibco 15140). 293T cells were grown in Dulbecco’s Modified Eagle Medium (DMEM, Gibco 11995) supplemented with 10% FBS and 1% Pen Strep. Cells were cultured at 37 °C and 5% CO_2_.

### Generation of *PCNA^K164R^*, *RAD18^-/-^, RAD18^-/-^PCNA^K164R^* cell lines using CRISPR-Cas9

A guide RNA (gRNA) targeting PCNA exon 5 was designed (gRNA: 5’-ATACGTGCAAATTCACCAGA-3’) and cloned into a CRISPR/Cas9 (<u>c</u>lustered regularly interspaced <u>s</u>hort <u>p</u>alindromic repeats/<u>C</u>RISPR <u>a</u>ssociated <u>9</u>) plasmid (hSpCas9(BB)-2A-GFP/PX458; Addgene plasmid #48138) as described previously (Ran et al., 2013). To generate a *PCNA^K164R^* mutant cell line, a double stranded donor plasmid containing the desired K164R mutation was constructed using Golden Gate cloning and designed as described previously (Fattah et al., 2014; Kohli et al., 2004; Oh et al., 2014; Thompson et al., 2017). Silent mutations were introduced into the donor plasmid to generate a novel restriction enzyme recognition site. RPE-1 wildtype cells were transfected with the CRISPR/Cas9 plasmid containing the PCNA gRNA and the donor plasmid containing the K164R mutation using the Neon Transfection System (Invitrogen MPK5000) following standard protocols. Two days post-transfection GFP-expressing cells were collected by flow cytometry and subcloned. Subclones were screen for correct targeting by PCR amplification and restriction enzyme digestion (Forward: 5’-TGGCGCTAGTATTTGAAGCA-3’, Reverse: 5’-ACTTGGGATCCAATTCTGTCTACT-3’, Restriction Enzyme: EcoRI, NEB R3101). Specific mutations were identified by Sanger sequencing (Sequencing: 5’-AGGTGTTGCCTTTTAAGAAAGTGAGG-3’).

To generate *RAD18^-/-^* and *RAD18^-/-^:PCNA^K164R^* mutant cell lines, a gRNA targeting RAD18 exon 2 (gRNA: 5’-AGACAATAGATGATTTGCTG-3’) was designed such that DNA cleavage would disrupt an endogenous restriction enzyme recognition site and was subsequently cloned into a CRISPR/Cas9 plasmid as described above. RPE-1 wildtype cells and *PCNA^K164R^* 2B10 cells were transfected with the CRISPR/Cas9 plasmid containing the RAD18 gRNA using the Neon Transfection System following standard protocols. Two days post-transfection GFP-expressing cells were collected by flow cytometry and subcloned. Subclones were screen for correct targeting by PCR amplification and restriction enzyme digestion (Forward: 5’-GTAGTACCATGCCGAAAGCAC-3’, Reverse: 5’-GGAACCACCTATCTGTTATCC-3’, Restriction Enzyme: TseI, NEB R0591). Knockout lines were identified by Sanger sequencing (Sequencing: 5’-CTACCTCATGTAAAAATCGC-3’) and <u>T</u>racking of Indels by <u>DE</u>composition (TIDE) analyses (Brinkman et al., 2014). 293T *PCNA^K164R^* lines were generated as described previously (Thakar et al., 2020).

### Cell Proliferation

Cells were plated at 100,000 cells per well (RPE-1) or 125,000 cells per well (293T) in 6-well plates. Cell counts were performed 3-days after seeding using Trypan Blue (Invitrogen T10282) on Countess slides (Invitrogen C10283) using a Countess automated cell counter (Invitrogen C20181).

### Cell Viability Assay

RPE-1 cells were plated at 500 cells (wildtype) or 800 cells (*PCNA^K164R^*) per well in 96-well plates and allowed to recover for 24 h. Stock solutions of each drug were prepared in sterile 1X phosphate-buffered saline (PBS), water or dimethyl sulfoxide (DMSO) as appropriate and further diluted in growth medium. Cells were allowed to grow for 96 h in drug containing medium (methyl methanesulfonate, Acros Organics 156890250, mitomycin C, Sigma M4287, 4-nitroquinoline 1-oxide, Sigma N8141, hydroxyurea, Acros Organics 151680250; aphidicolin, Sigma A0781; camptothecin, Sigma C9911) and cell viability was measured with the CellTiter-Glo Luminescent Cell Viability Assay (Promega G7572) following manufacturer’s instructions. The viability of drug treated cells was normalized to the average viability of the untreated control cells for each cell line. Plates were imaged using a GloMax Discover Microplate Reader (Promega). Analysis and statistical test were performed using Microsoft Excel.

### Protein Extraction, Nuclear Fractionation and Western Blotting

For preparation of whole cell extracts, cells were lysed in NETN (20 mM Tris-HCl, pH 8.0, 100 mM NaCl, 1 mM EDTA, 0.5% NP-40 and protease inhibitors) buffer for 10 minutes at 4 °C and then centrifuged at 12,000 rpm for 10 minutes at 4 °C. Cleared lysates were collected and protein concentrations were determined using Bradford protein assay (Bio-Rad 5000006). Lysates were then mixed with SDS loading buffer and denatured at 95 °C before fractionation by SDS-PAGE and analyses by western blot. Nuclear fractions were isolated as previously described (Becker et al., 2018; Motegi et al., 2008). Briefly, extracts were prepared by lysis in Buffer A (10 mM HEPES pH 7.9, 10 mM KCl, 1.5 mM MgCl_2_, 0.34 M sucrose, 10% glycerol, 0.1% triton X-100 and protease inhibitors) for 5 minutes at 4 °C. Insoluble nuclear proteins were isolated by centrifugation at 1300 g at 4 °C and chromatin bound proteins were subsequently released by sonication after being resuspended in TSE buffer (20 mM Tris-HCl, pH 8.0, 500 mM NaCl, 2 mM EDTA, 0.1% SDS, 0.1% triton X-100 and protease inhibitors). Remaining insoluble factors were cleared by centrifugation at 17000 g at 4 °C. Protein concentrations of nuclear fractions were determined using Bradford protein assay prior to fractionation by SDS-PAGE and western blot analyses. Primary antibodies were incubated in 5% BLOT-QuickBlocker (G-Biosciences 786-011) as follows: mouse anti-PCNA (Abcam, ab29; 1:3000), rabbit anti-Ubiquityl-PCNA (Lys164) (Cell Signaling, D5C7P, 13439; 1:1000), rabbit anti-RPA32 (S33) (Bethyl, A300-246A; 1:2000), rabbit anti-γH2AX (Bethyl, A300-081A; 1:2000), rabbit anti-H2AX (Bethyl, A300-082A; 1:5000), rabbit anti-p-p53 (S15) (Cell Signaling, 9284S; 1:500), mouse anti-p53 (Santa Cruz, sc-126; 1:2000), rabbit anti-p21 (Santa Cruz, sc-397 clone C19; 1:1000), rabbit anti-FANCD2 (Abcam, ab108928; 1:2000), rabbit anti-RAD18 (Bethyl, A300-340A; 1:1000), rabbit anti-MCM2 (Cell Signaling, 4007S; 1:1000; BD Biosciences, 610701; 1:1000), rabbit anti-MCM3 (Cell Signaling, 4012S; 1:1000), rabbit anti-MCM4 (Cell Signaling, 3228S; 1:1000); rabbit anti-MCM7 (Cell Signaling, 3757S; 1:1000), mouse anti-GAPDH (GeneTex, GTX627408; 1:10000), mouse anti-Ku86 (Santa Cruz, B-1, sc-5280; 1:500), mouse anti-α-tubulin (Millipore, T9026, clone DM1A; 1:10000). Secondary antibodies were incubated in 5% BLOT-QuickBlocker as follows: goat anti-mouse HRP conjugate (BioRad, 1706516; 1:5000), donkey anti-rabbit HRP conjugate (Amersham, NA9340; 1:5000). Detection was performed using WesternBright Quantum detection kit (K-12042-D20). Quantification was performed using FIJI and Microsoft Excel. Image preparation was performed using Adobe Photoshop.

### FACS Analysis

For flow cytometry analyses of cell cycle distribution, DNA synthesis and origin licensing, wildtype and *PCNA^K164R^* RPE-1 and 293T cells were treated as described previously (Matson, eLife, 2017). Briefly, cells were treated with methyl methanesulfonate (20 µM) or mitomycin C (20 nM) for 48 h when applicable and incubated with 10 µM EdU (Lumiprobe, 20540) for 30 minutes before harvesting with trypsin. Cells were then washed with cold 1X PBS and lysed in CSK (10 mM PIPES pH 7.0, 300 mM sucrose, 100 mM NaCl, 3 mM MgCl_2_ hexahydrate) with 0.5% triton X-100, then fixed in PBS with 4% paraformaldehyde (Electron Microscopy Services) for 15 minutes. Cells were labeled with AF647-azide (Life Technologies, A10277) in 100 mM ascorbic acid, 1 mM CuSO_4_, and PBS to detect EdU for 30 minutes at room temperature (RT). Cells were then washed and incubated with anti-MCM2 antibody (BD Biosciences, #610700; 1:200) in 1% bovine albumin serum (BSA) in PBS with 0.1% NP-40 for 1 h at 37 °C. Next, cells were washed and labeled with donkey anti-mouse AF488 secondary antibody (Invitrogen, A11029; 1:1000) in 1% BSA in PBS with 0.1% NP-40 for 1 h at 37 °C. Lastly, cells were washed and incubated in DAPI (Life Technologies, D1306; 1 µg/mL) and RNAse A (Sigma, R6513; 100 ng/mL) for 1 h at 37 °C. Samples were processed on a LSR II (BD Biosciences) flow cytometer and analyzed with FlowJo v10.6.1 and Microscoft Excel.

For flow cytometry analyses of apoptosis, RPE-1 wildtype and *PCNA^K164R^* cells were seeded in 6-well plates at 75,000 cells per well and allowed to proliferate for approximately 72 h. Cells were collected, washed twice with 1X PBS and stained using the APC Annexin V apoptosis detection kit (Biolegend 640932) according to the manufacturer’s instructions. Samples were processed on a LSRII (BD Biosciences) flow cytometer. Apoptotic cells were identified by annexin V staining while cell viability was determined by PI staining. Data was analyzed using FlowJo v10.6.1 and GraphPad Prism 8.

### DNA combing

For genome-wide analyses of DNA replication, wildtype and *PCNA^K164R^* RPE-1 and 293T cells were plated at 40% confluency in 10 cm plates 24 h prior to labeling. Cells were incubated with 25 or 100 µM IdU (Sigma C6891) for 30 minutes, rinsed twice with pre-warmed medium and then incubated with 100 µM or 200 µM CldU (Sigma I7125) for 30 minutes. Approximately 250,000 cells were embedded in 0.5% agarose plugs (NuSieve GTG Agarose, Lonza, 50080) and digested for 48 h in plug digestion solution (10 mM Tris-HCl, pH 7.5, 1% Sarkosyl, 50 mM EDTA and 2 mg/mL Proteinase K). Plugs were then melted in 50 mM MES pH 5.7 (Calbiochem #475893) and digested overnight with β-agarase (NEB M0392). DNA was then subsequently combed onto commercially available vinyl silane-coated coverslips (Genomic Vision COV-001). Integrity of combed DNA for all samples was quality checked via staining with YOYO-1 (Invitrogen Y3601). Combed coverslips were baked at 60 °C for 2-4 h, cooled to RT and stored at −20 °C. DNA was denatured in 0.5 M NaOH and 1 M NaCl for 8 minutes at RT. All antibody staining was performed in 2% BSA in PBS-Triton (0.1%). Primary antibodies included rabbit anti-ssDNA (IBL 18731), mouse anti-BrdU/IdU (BD Biosciences 347580; clone B44) and rat anti-BrdU/CldU (Abcam, ab6326; BU1/75 (ICR1)). Secondary antibodies included goat anti-mouse Cy3.5 (Abcam ab6946), goat anti-rat Cy5 (Abcam ab6565) and goat anti-rabbit BV480 (BD Horizon #564879). Imaging was performed using Genomic Vision EasyScan service. Images were blinded and analyzed using the Genomic Vision FiberStudio software. Data/statistical analyses were performed in Microsoft Excel and GraphPad Prism 8.

### Small-interfering RNA (siRNA) transfection

RPE-1 cells were seeded on coverslips (Thermo Fisher 3405) in 6-well plates and allowed to recover for 24 h. Cells were treated with MCM4 siRNA using Lipofectamine RNAiMAX (Thermo Fisher 13778) in Opti-MEM (Thermo Fisher 31985062) supplemented with 3% FBS for 48 h.

### Immunostaining

For immunofluorescent staining of 53BP1 nuclear bodies and Cyclin A, wildtype and *PCNA^K164R^* RPE-1 and 293T cells were seeded onto fibronectin (Sigma F4759) coated coverslips at 100,000 cells per coverslip and allowed to recover for 24 h. Cells were then treated with 300 nM APH (Sigma A0781) for 24 h. After, cells were washed with PBS containing 0.9 mM CaCl_2_ and 0.49 mM MgCl_2_ (PBS-Ca^2+^/Mg^2+^) and fixed in PBS-Ca^2+^/Mg^2+^ with 3.7% formaldehyde (Fisher Scientific F79-500) for 10 minutes at RT. Next, cells were washed twice with PBS-Ca^2+^/Mg^2+^, permeabilized with 0.1% triton X-100 for 5 minutes at RT, and subsequently blocked in ABDIL (20 mM Tris-HCl, pH 7.5, 150 mM NaCl, 2% BSA, 0.2% Fish Gelatin, 0.1% NaN_3_) for 1 h at RT. Coverslips were incubated with rabbit anti-53BP1 (Abcam, ab36823; 1:500) and mouse anti-Cyclin-A (Santa Cruz; sc-271682 clone B8; 1:200) primary antibodies overnight at 4 °C. Next day, coverslips were washed with PBS-Ca^2+^/Mg^2+^ containing 0.1% Tween 20 and incubated with Alexa Fluor 488 donkey anti-rabbit (Invitrogen, A21206; 1:1000) and Alexa Fluor 594 goat anti-mouse (Invitrogen, A11032; 1:000) secondary antibodies for 1 h at RT. Lastly, coverslips were washed, stained with DAPI (Life Technologies, D1306; 1 µg/mL), and mounted on microscope slides with Vectorshield anti-fade reagent (Vector Laboratories H1000). 30-40 fields per coverslip were imaged on the Zeiss Spinning Disc confocal (University of Minnesota Imaging Center). Images were scored using FIJI and statistical analyses were performed in GraphPad Prism 8.

For immunofluorescent staining of FANCD2 and EdU foci, wildtype and *PCNA^K164R^* RPE-1 and 293T lines were seeded onto fibronectin coated coverslips and treated with 300 or 450 nM APH when applicable. Cells were then washed with PBS and incubated with 20 µM EdU for 30 minutes before fixation with 10% formalin. Fixed cells on coverslips were then subjected to a Click-Chemistry Reaction (20 µM Biotin-Azide, 10 mM sodium ascorbate, and 2 mM CuSO_4_ in PBS) at RT for 1 h. After, coverslips were washed with PBS and incubated with rabbit anti-FANCD2 (Abcam, ab108928; 1:250) and mouse anti-phospho-Histone H3 (S10) (Cell Signaling; 9706S; 1:200) primary antibodies in PBS containing 0.3% triton X-100 and 1% BSA at 4 °C overnight. The next day, coverslips were washed with PBS and incubated with Alexa Fluor 488 Streptavidin (Thermo Fisher S32354; 1:100), Alexa Fluor 350 anti-mouse (Thermo Fisher A11045; 1:100) and Alexa Fluor anti-rabbit (Thermo Fisher A31632; 1:1000) secondary antibodies at RT for 1 h. Lastly, coverslips were washed with PBS and mounted on microscope slides with Prolong Gold anti-fade reagent (Thermo Fisher P36931). EdU and FANCD2 foci were scored using a Zeiss Axio Imager A1 fluorescent microscope and EVOS FL imaging system (ThermoFisher AMF43000). 300-400 cells per cell line were scored per experiment. Statistical analyses were performed in GraphPad Prism 8.

For immunofluorescent staining of anaphases, wildtype and *PCNA^K164R^* RPE-1 cells were seeded onto coverslips at 200,000 cells per coverslip and allowed to recover for 24 h. Cells were then treated with 300 nM APH for 24 h. After, cells were washed with PBS and fixed with 10% formalin for 15 minutes. Fixed cells on coverslips were then washed with PBS, stained with DAPI and mounted on microscope slides with Vectorshield anti-fade reagent. 200-300 anaphases per cell line were scored per experiment using the EVOS FL imaging system (ThermoFisher AMF43000). Statistical analyses were performed in GraphPad Prism 8.

## ACKNOWLEDGEMENTS

We would like to thank the University of Minnesota Flow Cytometry Core Facility, Imaging Center and Cytogenomics Shared Resource.

## AUTHOR CONTRIBUTIONS

Conceptualization, A.K.B. and W.L.; Methodology, A.K.B., W.L., N.S.; Formal Analysis, A.K.B., W.L.; Investigation, W.L., R.M.B., T.T., Y.C.C., L.W., C.B.R., W.D., A.T.; Writing – Original Draft, W.L., and A.K.B.; Writing – Review & Editing, W.L., R.M.B., A.K.B., G.L.M., N.S.; Visualization, W.L., Supervision, A.K.B., Project Administration, W.L. and A.K.B.; Funding Acquisition, A.K.B.

---

**Figure S1.**
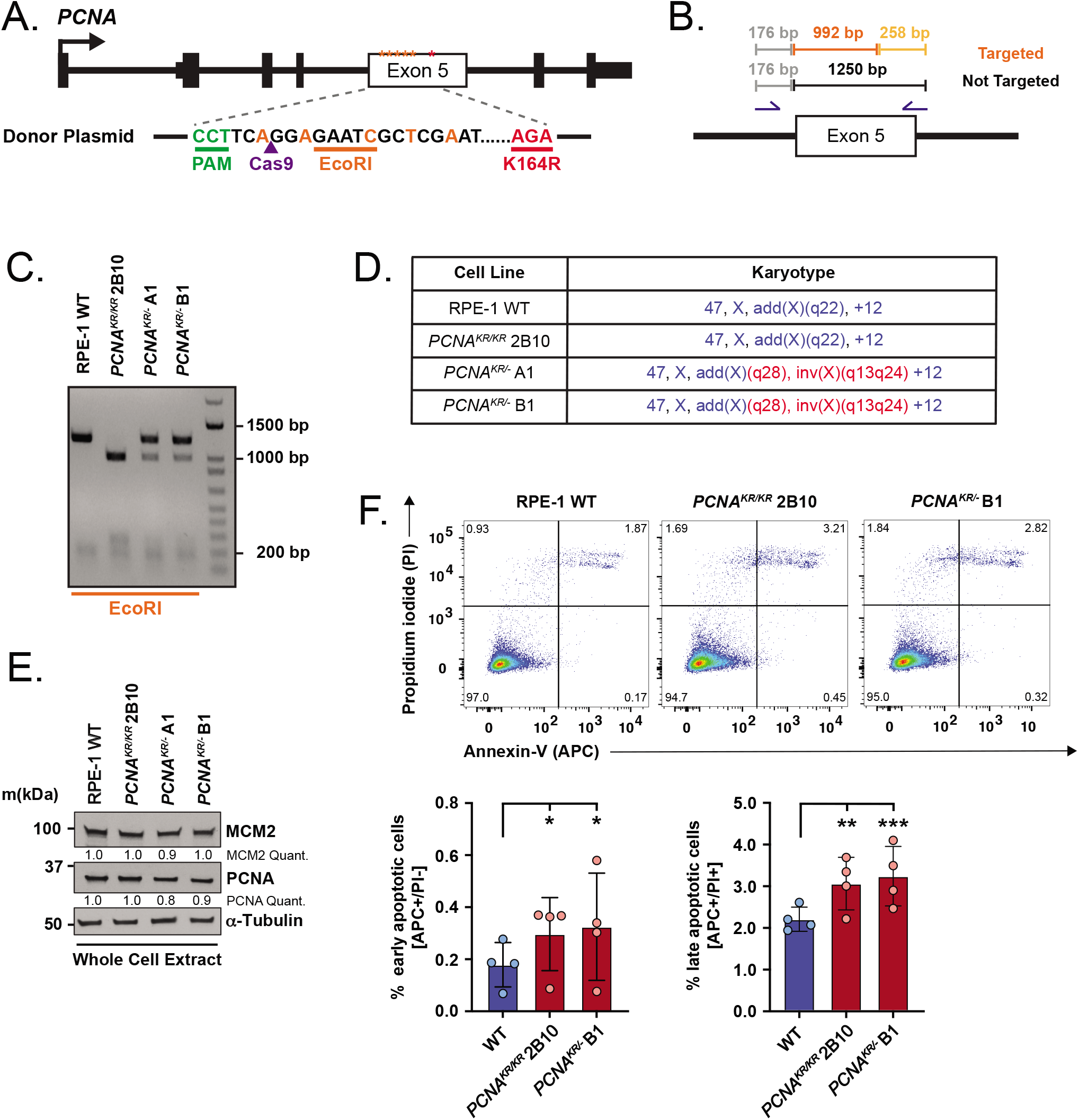
Generation of a *PCNA^K164R^* mutant cell line in RPE-1 using CRISPR/Cas9. **A)** Schematic of the human *PCNA* indicating that exon 5 was targeted by CRISPR-Cas9. The K164R mutation was knocked-in utilizing a donor plasmid. **B)** Schematic of screening PCR and expected PCR product sizes after EcoRI restriction enzyme digestion. **C)** Representative genotyping PCR. Not targeted (wildtype; 176bp, 1250 bp), monoallelic knock-in (KIN) (*PCNA^KR/-^* A1 and B1; 176 bp, 258 bp, 992 bp, 1250 bp), and biallelic KIN (*PCNA^KR/KR^* 2B10; 176 bp, 258 bp, 992 bp). **D)** Karyotype analysis of RPE-1 wildtype, *PCNA^KR^* (2B10) and *PCNA^KR/-^* (A1 and B1) cell lines. Blue indicates expected RPE-1 karyotype. Red indicates chromosomal abnormalities. **E)** Western blot analyses of whole cell extracts from wildtype RPE-1 and *PCNA^K164R^* cells for MCM2 and PCNA with α-tubulin as the loading control. Intensities of MCM2 and PCNA levels normalized to loading control. **F)** (Top) Representative flow cytometry plots sorting cells based on propidium iodide and annexin V staining for the quantification of early apoptotic, late apoptotic and dead cells in RPE-1 wildtype and *PCNA^K164R^* cell lines. (Bottom) Percentage of early apoptotic (left) and late apoptotic (right) cells. Error bars indicate standard deviation and significance was calculated using students *t-test* with *>.05; **>.01, ***>.001; n=12 replicates across four biological replicates.

**Figure S2.**
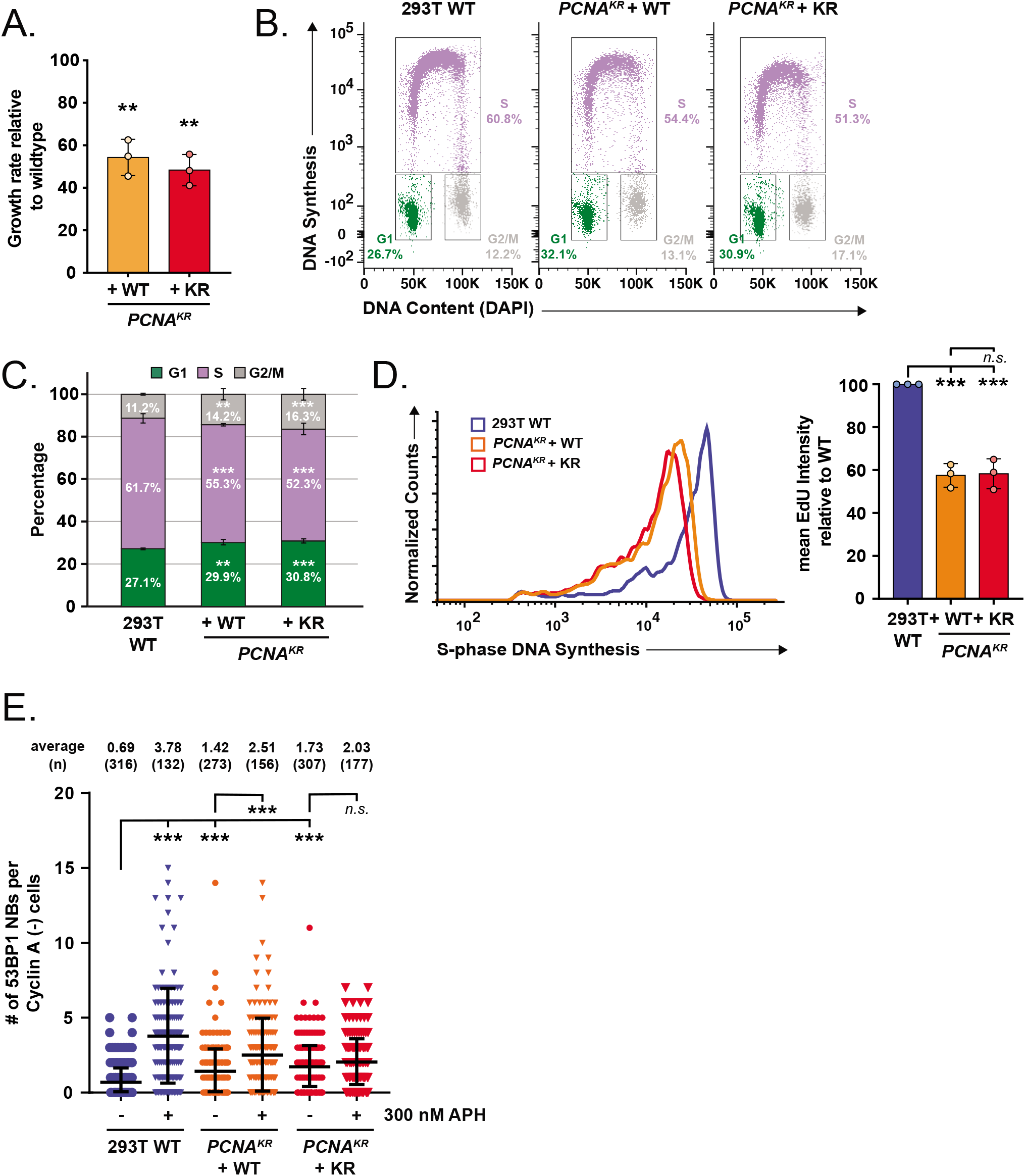
293T *PCNA^K164R^* mutant cells exhibit significant DNA synthesis defects. **A)** Average cell proliferation rate in 293T *PCNA^K164R^* lines complemented with either wildtype or K164R cDNA normalized to wildtype. For each cell line, n = 9 wells across three biological replicates. Error bars indicate standard deviation and significance was calculated using students *t-test* with **>.01. **B)** Representative cell cycle distribution of 293T wildtype and *PCNA^K164R^* lines complemented with either wildtype or K164R cDNA based on DNA content (DAPI) and DNA synthesis (EdU incorporation). Percentage of each population in G1- (green), S- (purple) or G2/M-phase (gray) is shown. **C)** Cell cycle distribution of 293T wildtype and *PCNA^K164R^* lines complemented with either wildtype or K164R cDNA from three biological replicates. Percentage of each population in G1- (green), S- (purple) or G2/M-phase (gray) is shown. Error bars indicate standard deviation and significance was calculated using students *t-test* with ***>.001. Please note that EdU incorporation reported by Thakar et al., 2020 was performed using different experimental conditions. **D)** Histogram (left) and quantification of mean fluorescent intensity (right) of EdU staining of S-phase cells from 293T wildtype (blue) and *PCNA^K164R^* lines complemented with either wildtype (orange) or K164R (red) cDNA, n = 9 across three biological replicates. Error bars indicate standard deviation and significance was calculated using students *t-test* with *>.05. **E)** 53BP1 NB quantification of untreated (circles) and APH treated (triangles) 293T wildtype (blue) and *PCNA^K164R^* lines complemented with either wildtype (orange) or K164R (red) cDNA. Number (n) of nuclei quantified is listed. Error bars indicate standard deviation and significance was calculated using one-way ANOVA with Tukey’s multiple comparison test with ***<.001.

**Figure S3.**
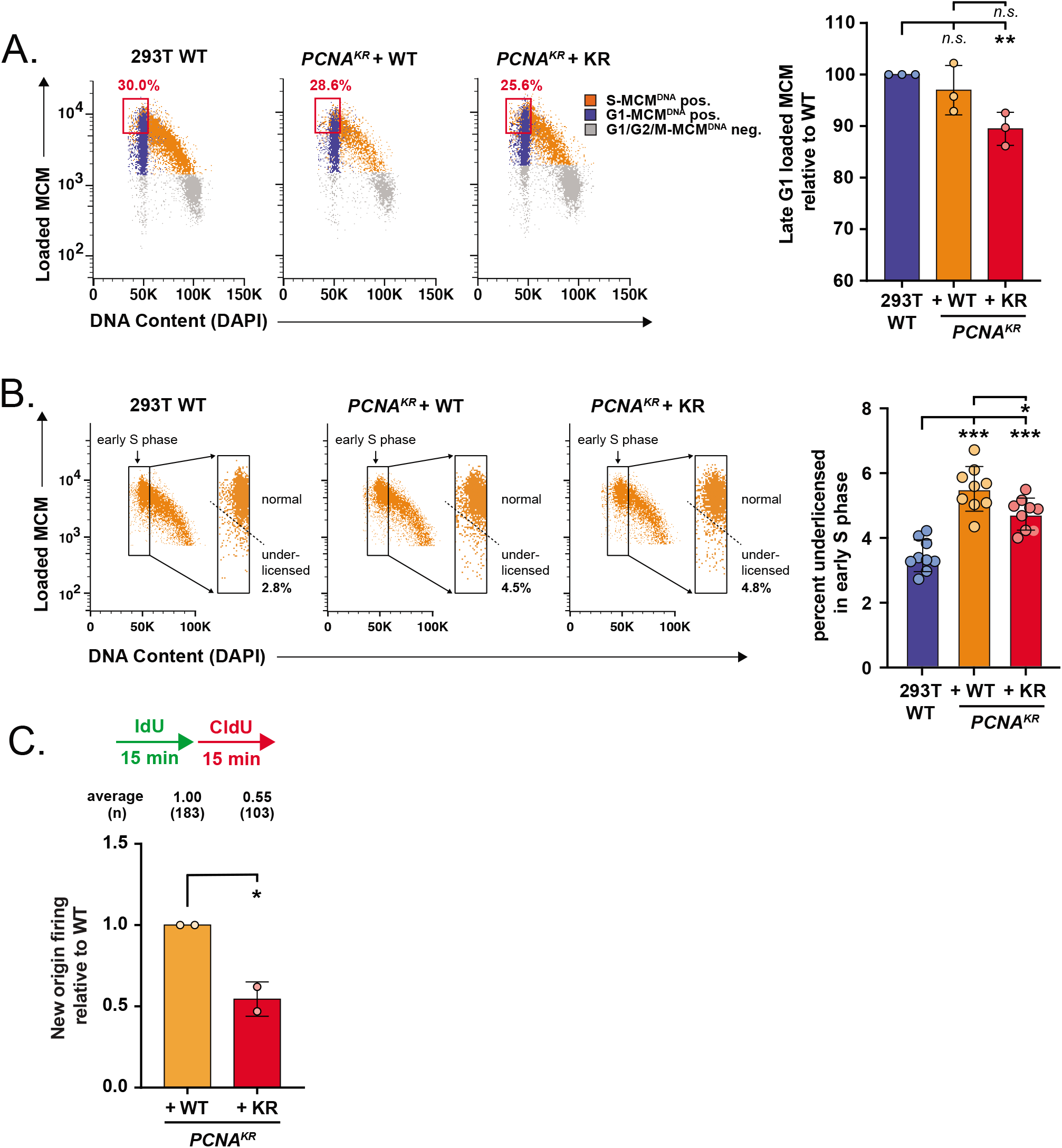
Reduced MCM2 loading in 293T *PCNA^K164R^* cells. **A)** (Left) Representative chromatin flow cytometry plots for 293T wildtype, *PCNA^K164R^* cells complemented with either wildtype PCNA or a K164R cDNA. G1 phase/MCM positive cells (blue), S phase/MCM positive cells (orange) and G1- or G2/M-phase/MCM negative cells (gray) are indicated. Percentage of MCM2 stained cells in late G1 is indicated (red box). (Right) Quantification of the percentage of MCM2 stained cells in late G1 from 293T wildtype (blue) and *PCNA^K164R^* cells complemented with wildtype (orange) or a K164R (red) cDNA; n = 9 across three biological replicates. Error bars indicate standard deviation and significance was calculated using students *t-test* with *>.05. **B)** (Left) Gating of early S phase cells from A), showing only the S phase, MCM/DNA-positive cells. Rectangles define early S phase cells as MCM/DNA-positive with 2C DNA content. The dashed line defines the cutoff between normally licensed versus under-licensed cells. (Right) Percentage of under-licensed cells in early S phase from 293T wildtype (blue) and *PCNA^K164R^* cells complemented with wildtype (orange) or a K164R (red) cDNA; n = 9 across three biological replicates. Error bars indicate standard deviation and significance was calculated using students *t-test* with *>.05, **>.01, ***>.001. **C)** Active replication forks were sequentially labeled with IdU (100 µM, green) for 15 minutes followed by labeling with CldU (100 µM, red) for 15 minutes. NOF is measured as CldU-only tracts. NOF events under unperturbed conditions from 2 biological replicates in 293T *PCNA^K164R^* lines complemented with either wildtype (orange) or K164R (red) cDNA normalized to wildtype. Number (n) of events quantified is listed. Significance was calculated using students *t-test* with *>.05.

**Figure S4.**
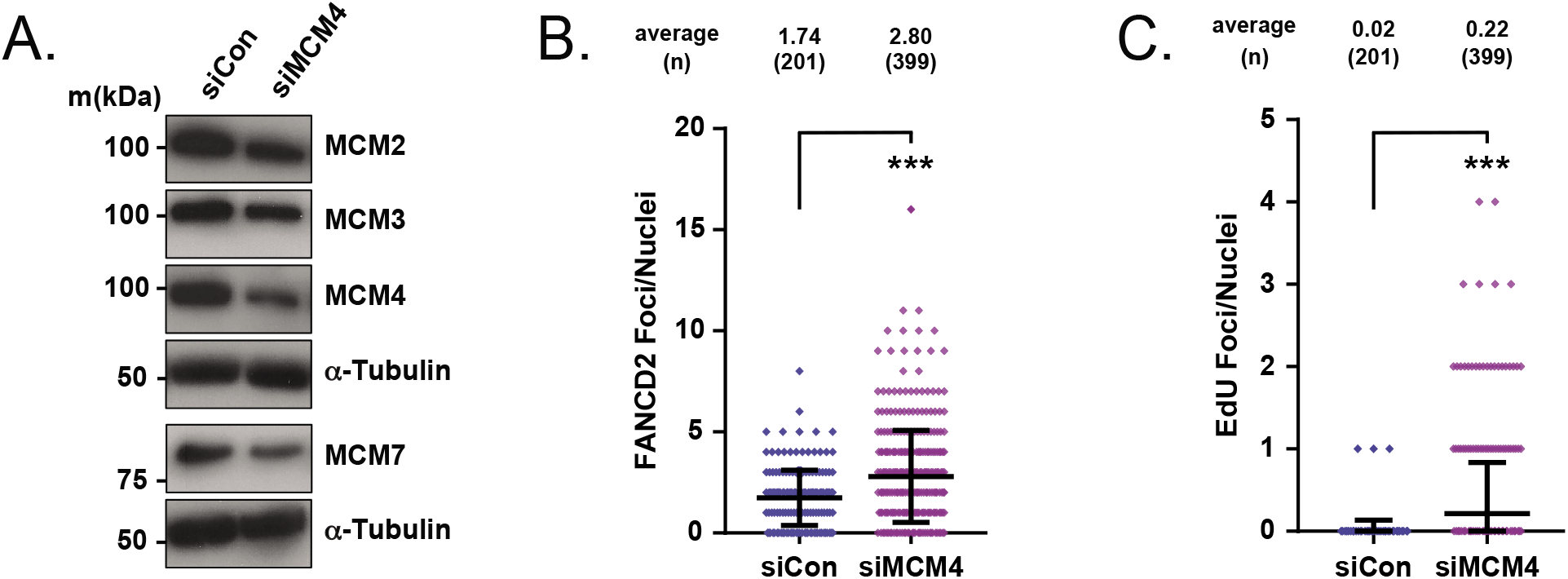
Enhanced MiDAS activation in MCM4 knockdown cells. **A)** Western blot analyses of whole cell extracts from wildtype RPE-1 cells treated with siControl or siMCM4 for MCM2, MCM3, MCM4, and MCM7 with α-tubulin as the loading control. **B)** FANCD2 foci quantification of two biological replicates in wildtype RPE-1 cells treated with siControl (blue) or siMCM4 (purple). Number (n) of nuclei quantified is listed. Significance was calculated by students *t-test* with ***<.001. **C)** EdU foci quantification of two biological replicates in wildtype RPE-1 cells treated with siControl (blue) or siMCM4 (purple). Number (n) of nuclei quantified is listed. Significance was calculated by students *t-test* with ***<.001.

**Figure S5.**
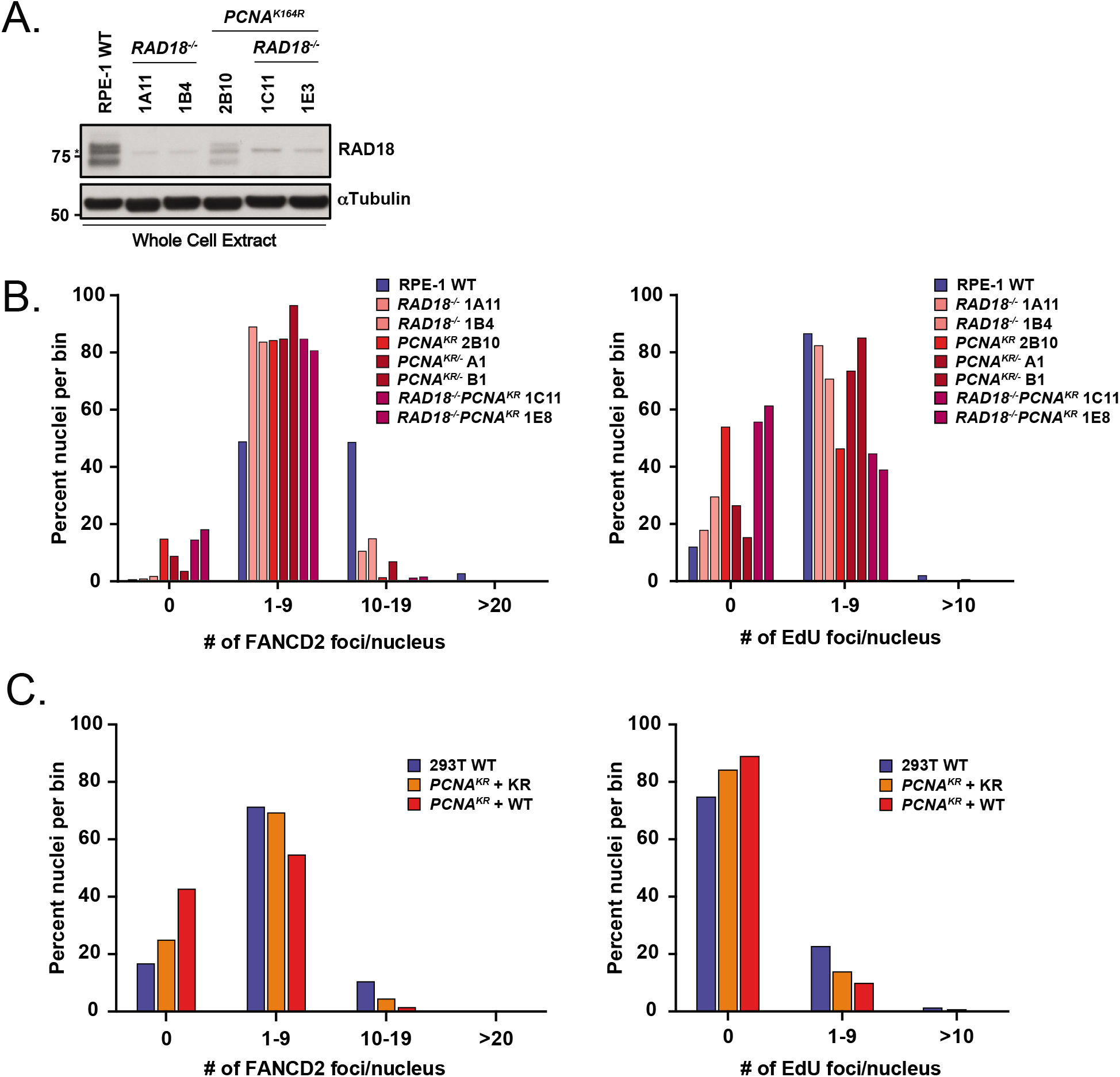
Reduced FANCD2 and EdU foci in *PCNA^K164R^* mutants. **A)** Western blot analyses of whole cell extracts from wildtype, *RAD18^-/-^*, *PCNA^K164R^* and *RAD18^-/-^PCNA^K164R^* RPE-1 cells for RAD18 with tubulin as the loading control. **B)** Bins of 0, 1-9, 10-19 and >20 FANCD2 foci and EdU foci per nucleus from two biological replicates of RPE-1 wildtype (blue), *RAD18^-/-^* (pink), *PCNA^K164R^* (red), *PCNA^K164R/-^* (maroon) and *RAD18^-/-^PCNA^K164R^* (purple) cells treated with APH. **C)** Bins of 0, 1-9, 10-19 and >20 FANCD2 foci and EdU foci per nucleus from two biological replicates in 293T wildtype (blue) and *PCNA^K164R^* cells complemented with either wildtype (orange) or a K164R (red) cDNA treated with APH.

